# DancE: A DNA-Activated Bacterial cGAS-like Enzyme in Antiphage Immune Defense

**DOI:** 10.64898/2026.10.06.757031

**Authors:** Yan Liu, Menghang Huang, Justin Jenson, Diana R. Tomchick, Zhijian J. Chen

**Affiliations:** Department of Molecular Biology; Center for Inflammation Research; Departments of Biophysics and Biochemistry University of Texas Southwestern Medical Center; Current address: Department of Biochemistry and Biophysics University of North Carolina at Chapel Hill; Howard Hughes Medical Institute

## Abstract

Vertebrate cGAS senses DNA and synthesizes cyclic GMP–AMP to launch an immune response through the adaptor protein STING. Bacterial cyclic oligonucleotide-based signaling systems (CBASS) are evolutionarily related to the metazoan cGAS–STING pathway, but the molecular cues triggering bacterial cGAS activation remain poorly understood. Here, we identify a cGAS-like enzyme, termed DancE (<u>D</u>NA-<u>a</u>ctivated di- <u>n</u>ucleotide <u>c</u>yclase in *<u>E</u>scherichia*), that is activated by binding to double-stranded DNA. DancE is repressed by the bacterial chaperone DnaK, which prevents DNA binding by DancE. We further identified lambda phage prohead protease gpC that activates DancE through its protease activity. These results provide an example of a bacterial cGAS-like enzyme that is activated by DNA but through a guard-like mechanism that normally suppresses DNA sensing, suggesting that DNA sensing by cGAS family proteins is an ancient antiviral strategy conserved across kingdoms of life.

---

Recent studies have revealed that bacterial antiphage defense systems are evolutionarily related to metazoan innate immune pathways (*1*). One prominent example is bacterial CBASS, which was reported to be found in 13% of investigated genomes (*2*). CBASS systems are classified into four types, of which Type I is the simplest, comprising a single cGAS/DncV-like cyclic dinucleotide synthase (CD-NTase) and an effector protein (*2*). Upon sensing phage infection, CD-NTases synthesize cyclic nucleotide second messengers that activate downstream effectors (*3*), leading to bacterial growth arrest or cell death to block phage propagation (*4*). Despite this well-established framework, the mechanisms by which bacterial CD-NTases are activated during phage infection remain poorly understood. Unlike mammalian cGAS, which is robustly activated by DNA both *in vitro* and *in vivo* (*5, 6*), several bacterial CD-NTases display little or no activity *in vitro*, suggesting the requirement for activating ligands (*3, 7*). Conversely, other CD-NTases, such as DncV, exhibit constitutive activity *in vitro*, implying the existence of host-encoded inhibitory factors that restrain signaling in the absence of infection (*3, 8*). Here, we show that the CD-NTase from *Escherichia albertii* is directly activated by DNA and that this activation is inhibited by the bacterial Hsp70 chaperone DnaK *in vitro*. We name this enzyme DancE (<u>D</u>NA-<u>a</u>ctivated di-<u>n</u>ucleotide <u>c</u>yclase in *<u>E</u>scherichia*). We further demonstrate that the lambda phage prohead protease gpC activates a DancE gain-of-function mutant *in vivo*, and that the activation depends on DNA binding by DancE. Together, these findings reveal a regulated-DNA sensing mechanism in which host and phage factors control CBASS activation during phage infection.

## Identification of DancE

Unlike mammalian cGAS that is known to produce 2’3’-cGAMP, bacterial cGAS-like enzymes from different species can produce a variety of cyclic di-nucleotides and cyclic oligonucleotides (*3*), raising the question of whether different cyclic di-nucleotides besides 2’3’-cGAMP exist in mammalian cells. If so, do they have functions beyond activating the STING pathway? To address this question, we leveraged the diversity of bacterial cGAS homolog CD-NTases that synthesize a broad repertoire of cyclic nucleotides (*3*). We selected a panel of CD-NTases that were reported to exhibit detectable enzymatic activity *in vitro* to enable intracellular production of diverse cyclic nucleotides (*2–4, 9*). Sixteen selected CD-NTases were transiently expressed in human HEK 293T cells, which lack STING expression, and their effects on cellular morphology and viability were monitored (Fig. 1A and table S1). This screen identified CD-NTase015 (*3*) (now renamed DancE) from *Escherichia albertii* as being capable of suppressing cell growth and inducing cell detachment from the culture plate (fig. S1). Subsequent mass spectrometry analysis showed that DancE predominantly produces cyclic di-adenosine monophosphate (c-di-AMP) (Fig. 1B and fig. S2A-S2C). The DancE system belongs to the Type I CBASS family and encodes a single CD-NTase and a two–transmembrane (2TM) effector protein, Cap15 (*3, 10, 11*) (Fig. 1C). DancE comprises an N-terminal catalytic nucleotidyltransferase domain (cyclase) and <u>A</u>denylyl/<u>G</u>uanylyl and <u>S</u>MODS <u>C</u>-terminal domain (AGS_C), a conserved feature of many Type I CBASS and some Type II CBASS (*2, 11*) (Fig. 1D). Cap15 contains an N-terminal transmembrane domain (TM) and a C-terminal nucleotide binding domain (NBD) (Fig. 1D). Binding of the cyclic nucleotide second messenger activates Cap15 to induce inner-membrane disruption and cell death, thereby mediating antiphage defense (*10*).

**Fig. 1.**
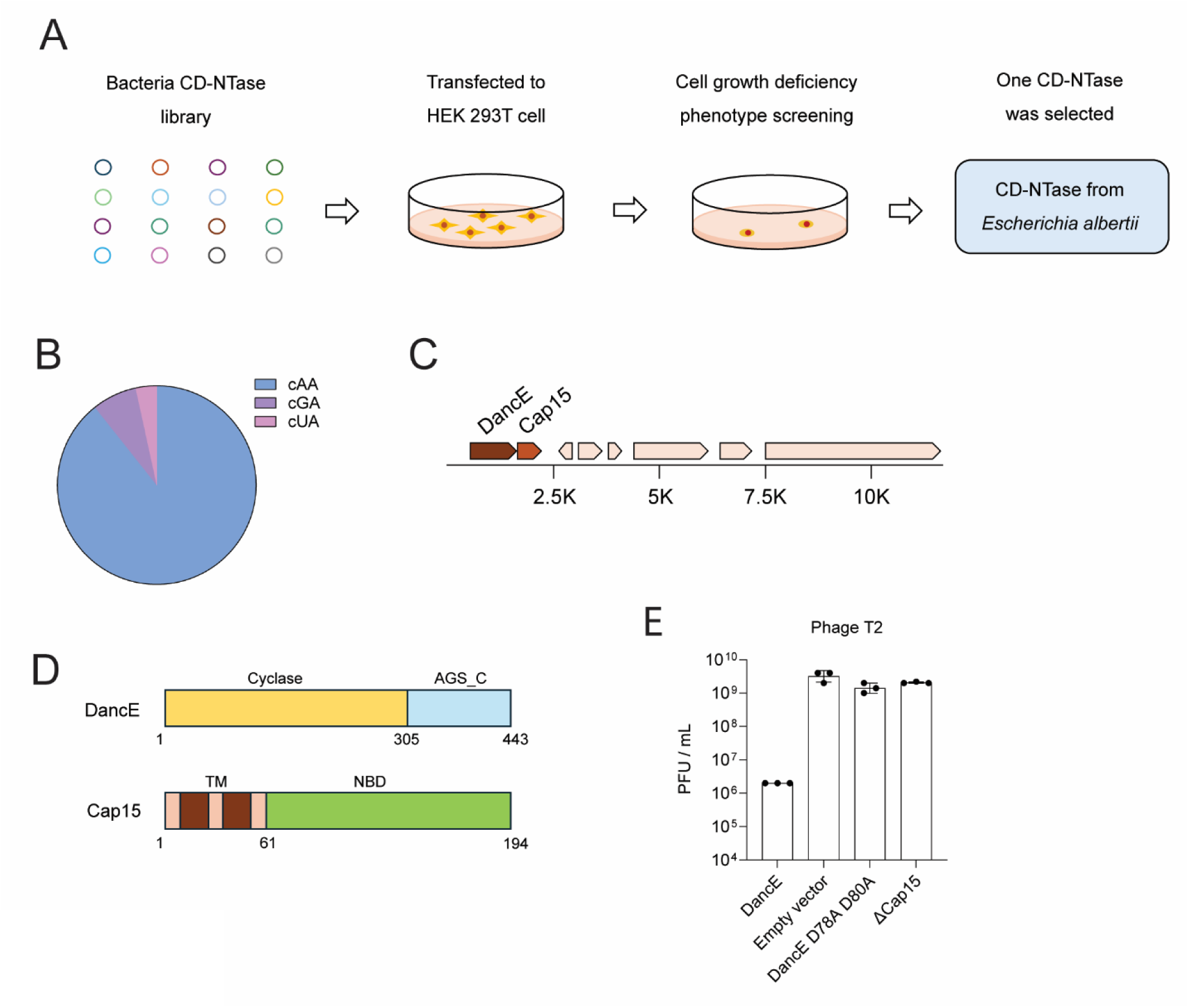
Identification of DancE. (**A**) Screening of CD-NTases for their ability to induce phenotypes in HEK 293T cells. CD-NTase from *Escherichia albertii* (DancE) was selected for inducing a significant cellular phenotype. (**B**) The major *in vivo* product of DancE in HEK 293T cells is c-di-AMP (cAA). Trace amounts of c-GMP-AMP (cGA) and c-UMP-AMP (cUA) were also detected. The data are representative of at least three independent experiments. (**C**) Domain organization of the CBASS operons from *Escherichia albertii*. (**D**) Schematic of DancE and Cap15 proteins. AGS_C, Adenylyl/Guanylyl and SMODS C-terminal sensor domain. TM, transmembrane domain. NBD, nucleotide binding domain. (**E**) Efficiency of plating (EOP) of T2 phage on *E. coli* MG1655 expressing the wild-type DancE CBASS operon, empty vector (no operon), D78A D80A mutant (DancE catalytic dead), or Cap15 deletion mutant of DancE CBASS. Data are mean ± s.d. from three biological replicates.

Our finding that overexpression of DancE but not its catalytic mutant (D78A D80A) causes cell detachment and morphology changes in HEK293T cells (fig. S1 and fig. S2C) suggests that cyclic di-AMP triggers a cellular response through a STING-independent mechanism. While we are still in the process of investigating this mechanism, our parallel studies of the role of DancE in bacterial antiphage immune defense led us to the unexpected discovery of DancE activation by double-stranded DNA and its regulation by phage infection, which are the subjects of this report.

## Structural Basis of DancE Architecture and Its Anti-Phage Function

To investigate the anti-phage function of DancE, we transformed the laboratory strain of *E. coli* MG1655, which does not harbor any known CBASS or CRISPR immune defense system, with the DancE CBASS operon. As reported previously (*11*), DancE CBASS conferred robust protection against T2 phage infection but did not inhibit infection by P1, lambda, T4, T5, or T6 phages (Fig. 1E and fig. S2D). The magnitude of protection was lower than that reported previously, likely reflecting the use of native promoter-driven expression in our study.

We determined the crystal structure of DancE at 2.55 Å resolution (Fig. 2A and table S2). The crystal structure revealed that DancE forms a tetramer organized as a dimer of dimers (Fig. 2A). Mass photometry analysis further supported the tetrameric assembly of DancE observed in the crystal structure (fig. S3A). In this arrangement, the spine regions of all four protomers are exposed on the outer surface of the assembly. In mammalian cGAS, the spine region mediates DNA binding (*12, 13*), suggesting that the corresponding surface of DancE may also participate in ligand recognition.

**Fig. 2.**
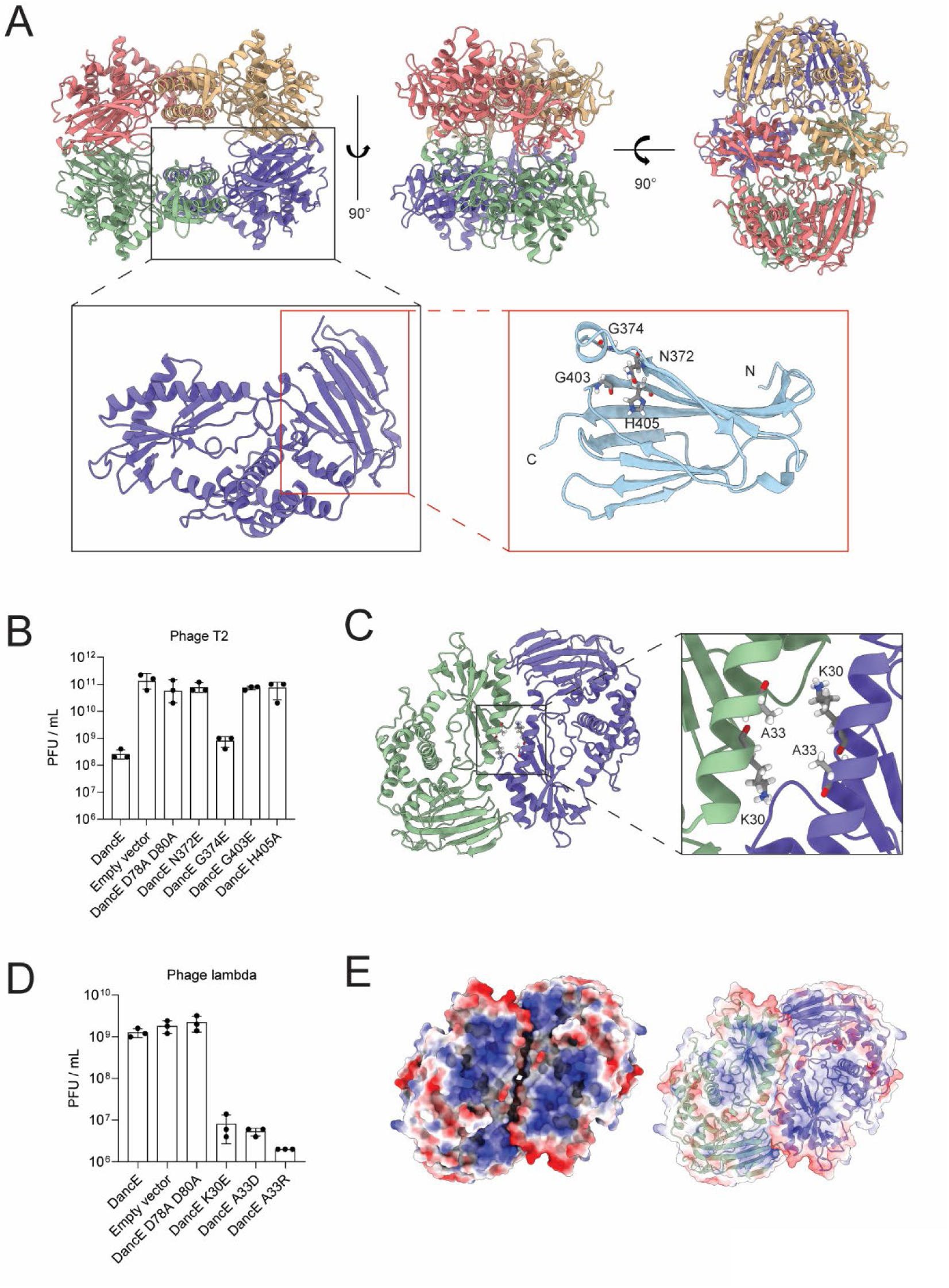
Structural Basis of DancE Architecture and Its Anti-Phage Function. (**A**) Crystal structure of apo DancE. Upper, DancE tetramer, different monomers are indicated by different colors. Lower left, DancE monomer. Lower right, DancE AGS_C domain, conserved sites studied in this study were indicated. N, N-terminal of AGS_C domain; C, C_terminal of AGS-C domain. Figures were created by ChimeraX. (**B**) Efficiency of plating (EOP) of T2 phage on *E. coli* MG1655 expressing wild-type operon, empty vector (no operon), D78A D80A mutant (DancE catalytic dead), or indicated AGS_C domain mutants of DancE CBASS. Data are mean ± s.d. from three biological replicates. (**C**) Left, crystal structure of DancE dimer. Right, enlarged dimer interface area. Figures were created by ChimeraX. (**D**) EOP of lambda phage on *E. coli* MG1655 expressing wild-type operon, empty vector, D78A D80A mutant, K30E, A33D, or A33R mutants of DancE CBASS. Data are mean ± s.d. from three biological replicates. (**E**) DancE dimer displayed with surface electrostatics (−10 to +10, red to blue) by ChimeraX.

Structural analysis of an individual protomer showed that DancE adopts the conserved CD-NTase fold, consisting of an N-terminal cyclase domain containing the catalytic center and spine region (Fig. 2A). This domain closely resembles mammalian cGAS. In addition, DancE contains a C-terminal AGS_C domain that forms a β-barrel-like structure. Previous studies have shown that the AGS_C domain exists in about 30% of type I CBASS CD-NTases and that this domain is important to enhance antiphage defense (*11*). Sequence analysis revealed several highly conserved residues within the AGS_C domain (Fig. 2A and fig. S3B). Mutation of the conserved sites (Asn372, Gly403, His405) resulted in loss of antiphage function (Fig 2B and fig. S3C), demonstrating that the AGS_C domain is essential for DancE function.

We also determined an independent crystal structure of DancE at 2.86 Å resolution using protein that had not undergone anion-exchange purification. This structure revealed a tetrahydrofolate (THFA) molecule bound in a surface pocket distal to the catalytic site. THFA binding caused only minor local conformational changes without altering the overall structure (fig. S3D and table S2). Folate derivatives have been reported to inhibit DncV (*8*). Consistent with the previous study, THFA inhibited the enzymatic activity of purified apo DancE, although relatively high concentrations of THFA were needed (fig. S3E). It is not clear whether folate derivatives may be involved in regulating bacterial CD-NTase activity *in vivo*.

To investigate the function of DancE oligomerization, we introduced mutations to Lys30 and Ala33 locating in DancE dimer interface to break the dimer (Fig. 2C). Surprisingly, rather than impairing antiphage defense, these mutations conferred a gain-of-function antiviral phenotype against lambda phage (Fig. 2D, fig. S3C and S3F). Substitution of Lys30 with glutamic acid (K30E) or Ala33 with either aspartic acid (A33D) or arginine (A33R) markedly enhanced resistance to lambda infection. Notably, lambda phage is not restricted by wild-type DancE (fig. S2D), suggesting that the dimer interface is involved in phage infection sensing and perturbation of the dimer interface may expand the antiviral specificity of DancE.

## DNA activates DancE to trigger anti-phage signaling

Structural analysis also revealed prominent positively charged surface regions on the DancE dimer opposite the catalytic pocket (Fig. 2E). The electrostatic properties of these patches suggest a potential role in engaging negatively charged ligands, such as nucleic acids. To investigate whether nucleic acids could function as activating ligands for DancE, we tested a panel of potential ligands, including double-stranded DNA (dsDNA), single-stranded DNA (ssDNA), double-stranded RNA (dsRNA) and single-stranded RNA (ssRNA), all at the same length (45 bp or nt). The dsRNA mimic poly(I:C) was also tested. Among these candidates, dsDNA elicited most robust activation of DancE, while ssDNA showed lower activation effect, and RNA or poly(I:C) showed no activation (Fig. 3A and fig. S4A), demonstrating that DancE is preferentially activated by dsDNA. Upon DNA stimulation, DancE predominantly generated c-di-AMP (accounted for approximately 75% of all detected cyclic nucleotides), as verified by chromatographic separation and mass spectrometry analysis (fig. S4B-S4D). In addition to c-di-AMP, low levels of other cyclic dinucleotides were also detected, indicating that DancE exhibits limited substrate promiscuity.

**Fig. 3.**
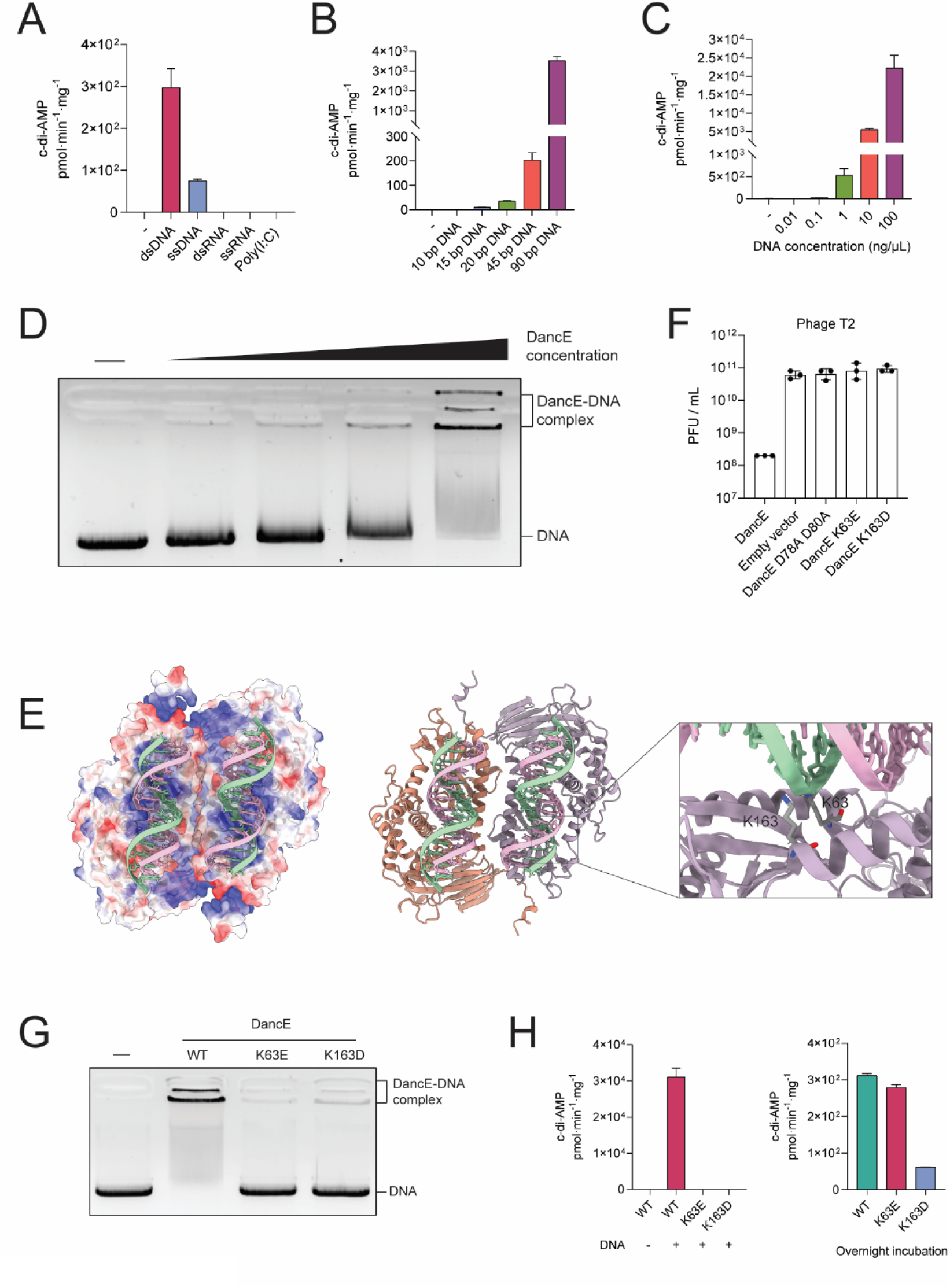
DNA activates DancE to trigger anti-phage signaling. (**A**)-(**C**) DancE is potently activated by DNA. (**A**) Wild-type DancE protein (0.8 µM) was incubated with or without indicated DNA or RNA or poly(I:C) (5 ng/µl) for 2 h. (**B**) Wild-type DancE protein (0.8 µM) was incubated with or without DNA of indicated length (10 ng/µl) for 2 h. (**C**) Wild-type DancE protein (0.8 µM) was incubated with or without indicated concentrations of 45-bp DNA for 2 h. Data are mean ± s.d. from three independent replicates. (**D**) DancE–DNA complex formation was assessed by EMSA using wild-type DancE protein at final concentrations of 0, 0.5, 1, 2, and 5 µM. (**E**) AlphaFold 3-predicted model of the DancE–DNA complex displayed with surface electrostatic potential (−10 to +10, red to blue) in ChimeraX. Inset, K63 and K163 within the predicted DNA-binding interface. ipTM = 0.81 and pTM = 0.86. (**F**) Efficiency of plating (EOP) of T2 phage on *E. coli* MG1655 expressing wild-type operon, empty vector (no operon), D78A D80A mutant (DancE catalytic dead) and DNA binding mutants of DancE CBASS. Data are mean ± s.d. from three biological replicates. (**G**) DancE–DNA complex formation was assessed by EMSA using wild-type or indicated mutant DancE proteins at final concentrations of 5 µM. (**H**) DNA binding mutations abrogate DNA activation of DancE. Left, wild-type or DNA binding mutant DancE proteins (0.8 µM) were incubated with or without plasmid (5 ng/µl) for 2 h. Right, wild-type or DNA binding mutant DancE proteins (3.5 µM) were incubated overnight (18 h). Data are mean ± s.d. from three independent replicates.

We then investigated the parameters governing DNA-dependent activation. At a fixed DNA concentration, longer DNA fragments induced significantly stronger activation than shorter fragments (Fig.3B and fig. S4E), which is consistent with human cGAS that is activated by long DNA more efficiently than short DNA (*14*). A linearized plasmid (5.4 kb) activated DancE to an extent similar to its circular counterpart and exhibited comparable activity to both phage genomic DNA and bacterial genomic DNA (fig. S4E). DNA titration experiments revealed clear concentration-dependent activation of DancE, with increasing DNA concentrations leading to progressively enhanced c-di-AMP production (Fig. 3C). Together, these results indicate that DancE activation is driven by both DNA length and abundance. We next used an electrophoretic mobility shift assay (EMSA) to examine the ability of DancE to interact with DNA *in vitro* using a linearized plasmid (Fig. 3D). In the presence of DNA, DancE exhibited electrophoretic mobility shift on the gel, indicating DNA binding.

To investigate the DNA-binding sites of DancE, we analyzed a model of DancE-DNA complex using AlphaFold 3 (Fig. 3E). The high-confidence model suggests that a DancE dimer interacts with two DNA superhelices through the positively charged surface located opposite the catalytic pocket. Guided by this prediction, we mutated Lys63 and Lys163, 2 residues positioned at the predicted DNA-binding interface, to glutamate or aspartate. Both mutations eliminated anti–T2 phage activity (Fig. 3F and fig. S4F). EMSA showed that the K63E and K163D mutants of DancE did not bind DNA anymore (Fig. 3G and fig. S4A). Measurement of c-di-AMP synthesis showed that the K63E and K163D mutants of DancE could not be activated by DNA but retained its residual basal enzymatic activity in the absence of DNA (Fig. 3H). Taken together, these data suggest that DNA binding is necessary to activate DancE and is essential for its anti-phage activity.

Since T2 phage can effectively activate DancE and the T2 genome has a special modification wherein cytosine is replaced by 5-hydroxymethylcytosine (hmC) (*15*), we next examined whether 5-hmC modification influences DancE activation. We compared the ability of the standard DNA (containing cytosine) and 5-hmC modified DNA to activate DancE and found that both DNA stimulated DancE to a similar extent (fig. S4G), indicating that 5-hmC is not a major determinant of enzymatic activation. Thus, DancE appears to respond primarily to the physical properties of double-stranded DNA. In contrast, the constitutively active CD-NTase DncV was not further stimulated by DNA (fig. S4H and S4I).

To investigate whether DancE homologs also bind DNA, we used AlphaFold 3 to predict the structures of DancE homolog–DNA complexes. High-confidence models were obtained for DancE homologs from *Escherichia*, *Pectobacterium*, and *Alteromonas* in complex with DNA (fig. S5A–S5D), suggesting that DNA binding, and potentially DNA-mediated activation, is a conserved feature across bacterial species.

To determine how DancE signaling is sensed by its cognate effector, we used AlphaFold 3 to predict a complex between the DancE-associated effector Cap15 and c-di-AMP, the major product of DancE. The predicted complex exhibited a relatively high confidence score (fig. S6A). To determine whether c-di-AMP is the second messenger produced by DancE that binds to Cap15 *in vivo*, we constructed an *E. coli* strain in which DancE was expressed from an arabinose-inducible promoter, whereas a Flag-tagged Cap15 variant lacking its transmembrane domains (to prevent cell death) was constitutively expressed (fig. S6B). Following DancE induction, Cap15-associated cyclic nucleotides were isolated by anti-Flag immunoprecipitation and analyzed by mass spectrometry. Nearly all recovered cyclic dinucleotides corresponded to c-di-AMP (fig. S6B), indicating that c-di-AMP is the predominant signaling molecule generated by DancE *in vivo* and directly recognized by Cap15. To validate the predicted interaction, we introduced mutations into residues predicted by AlphaFold 3 to mediate c-di-AMP binding (F94A, F137A and R176D) or sterically restrict access to the binding pocket (I82E) within Cap15. The I82E, F137A and R176D mutations markedly impaired c-di-AMP association with the effector (fig. S6C) and abolished resistance to T2 phage infection (fig. S6D). The F94A mutation did not significantly impair the binding of Cap15 to c-di-AMP but abolished its antiphage defense, suggesting that this residue may be involved in the effector function of Cap15.Together, these results establish c-di-AMP as the physiologically relevant second messenger produced by DancE that binds to the Cap15 effector *in vivo*.

## DnaK prevents DNA-dependent activation of DancE

In contrast to mammalian cGAS, which resides in the cytosol and is spatially separated from genomic and mitochondrial DNA by membrane compartmentalization under physiological conditions (*16*), bacterial CD-NTases lack such segregation and are therefore constitutively exposed to chromosomal DNA. This raises an important question as to how DancE avoids aberrant activation in a DNA-rich intracellular environment.

We observed that following purification by a heparin affinity column, the DNA-stimulated activity of DancE was markedly enhanced, whereas addition of the heparin column flow- through strongly repressed DNA-dependent activation (Fig. 4A and 4B), suggesting the presence of an inhibitory factor in the flow-through fraction. This inhibitory activity was resistant to heat treatment at 95°C for 10 min (fig. S7A).

**Fig. 4.**
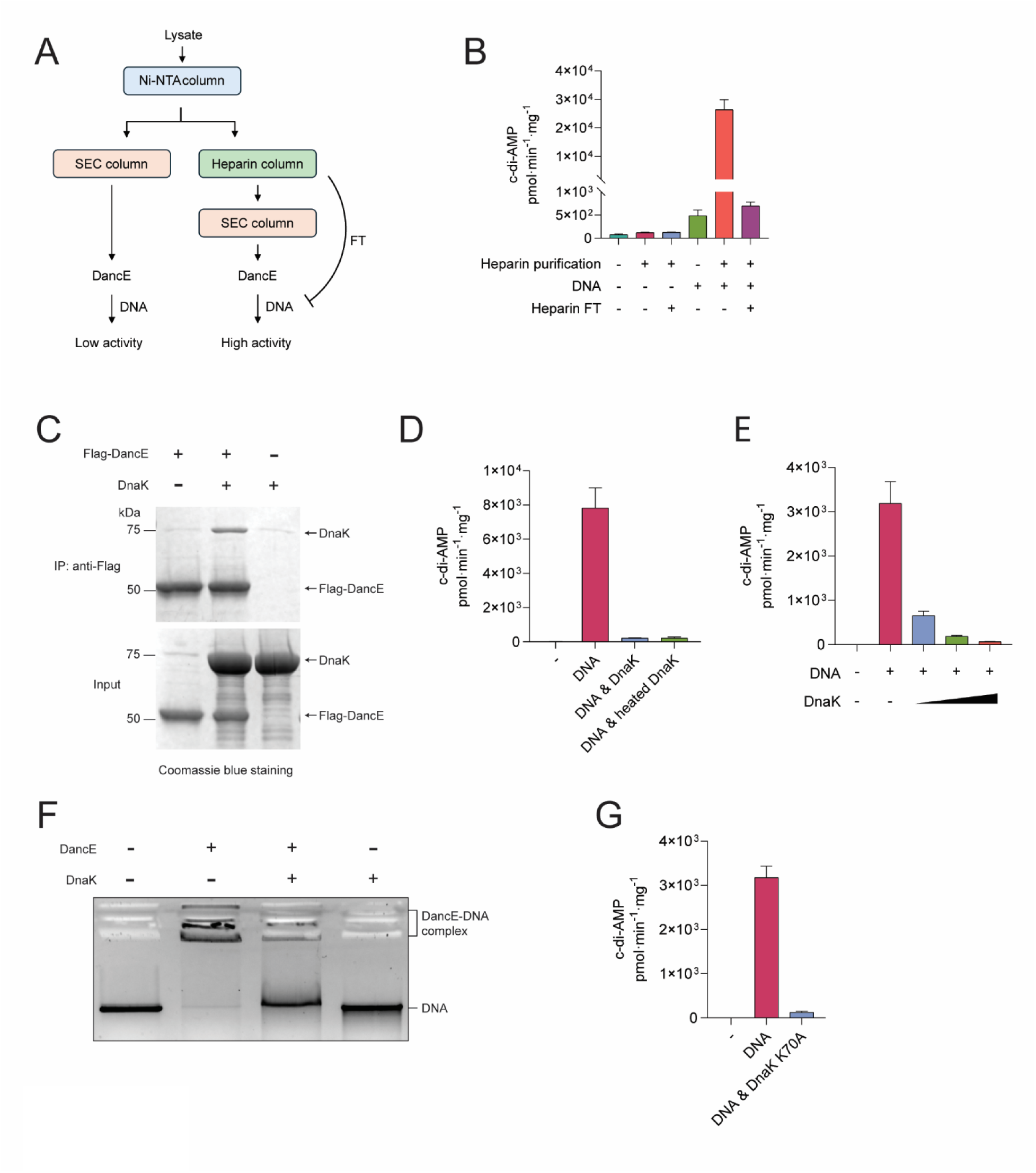
DnaK prevents DNA-dependent activation of DancE. (**A**) DancE purification workflow. The heparin column flow-through fraction represses DNA-induced activation of DancE. Ni–NTA, Nickel–nitrilotriacetic acid; SEC, size- exclusion chromatography. (**B**) Wild-type DancE protein (0.8 µM), purified with or without heparin column, was incubated with or without *E. coli* genomic DNA (5 ng/µl) and heparin column flow through (FT, 10% v/v) for 2 h. Data are mean ± s.d. from three independent replicates. (**C**) Immunoprecipitation analysis of DancE-DnaK interaction. Purified Flag– DancE and DnaK proteins were incubated with anti-Flag M2 magnetic beads (Sigma) for immunoprecipitation analysis. Samples were analyzed by SDS–PAGE followed by Coomassie blue staining. (**D**) Wild-type DancE (0.8 µM) was incubated with or without *E. coli* genomic DNA (5 ng/µl) in the presence or absence of native or heat-treated (95 °C for 10 min) DnaK (8 µM) for 1 h. Data are mean ± s.d. from three independent replicates. (**E**) Wild-type DancE (0.8 µM) was incubated with or without plasmid DNA (5 ng/µl) and increasing concentrations of DnaK (4, 8, or 16 µM) for 1 h. Data are mean ± s.d. from three independent replicates. (**F**) DancE–DNA complex formation was assessed by EMSA using wild-type DancE protein (5 µM final concentration), with or without DnaK protein (8 µM final concentration). (**G**) Wild-type DancE protein (0.8 µM) was incubated with or without plasmid DNA (5 ng/µl) and DnaK K70A protein (8 µM) for 1 h. Data are mean ± s.d. from three independent replicates.

To identify this factor, we further fractionated the heparin flow-through using an anion-exchange (Q) column and assessed the inhibitory activity of individual fractions. Mass spectrometry analyses of the fractions exhibiting robust repression of DNA-stimulated DancE activity identified DnaK as a protein whose abundance copurified with the inhibitory activity (fig. S7B). DnaK is a 70-kDa heat shock protein (Hsp70) that functions as a chaperone in protein folding and stress responses (*17*). Purified recombinant DnaK protein directly bound DancE and was sufficient to significantly repress DNA-induced activation of DancE *in vitro*, and this inhibitory effect remained heat resistant, confirming DnaK as the inhibitory factor present in the heparin flow-through fraction (Fig. 4C and 4D, and fig. S7C). The inhibitory effects on DancE activity increased with DnaK concentration, indicating a dose-dependent effect (Fig. 4E). EMSA further revealed that DnaK impaired DancE–DNA complex formation, suggesting that DnaK inhibits DancE activation by restricting DNA binding (Fig. 4F). To determine whether DnaK-mediated repression requires its ATPase activity, we introduced a K70A substitution that abolishes ATP hydrolysis. The DnaK K70A mutant retained the ability to suppress DNA-induced activation of DancE (Fig. 4G and fig. S7D), indicating that repression occurs independently of DnaK ATPase activity. DnaK also reduced the intrinsic enzymatic activity of DancE, albeit to a lesser extent, in the absence of DNA (fig. S7E).

Notably, DnaK did not repress the activity of DncV or human cGAS activation by DNA (fig. S7F-S7H), indicating that its inhibitory effect is specific to DancE rather than a general suppression of CD-NTase activity. Together, these results identify DnaK as a specific negative regulator of DancE, providing a mechanism by which DancE activity may be restrained in the presence of abundant genomic DNA.

## Lambda phage protease gpC triggers DancE activation

To investigate the mechanism of DancE activation by phage infection, we performed a forward genetic screen to isolate lambda phage escaper mutants capable of infecting bacteria expressing the gain-of-function DancE A33R variant, which restricts lambda phage infection (Fig. 2C and 2D). Using spontaneous mutagenesis and selection (Fig. 5A), we isolated two independent escaper phages that overcame DancEA33R–mediated restriction (Fig. 5B). Whole-genome sequencing revealed that both escapers harbored mutations in the lambda prohead protease gpC and the associated scaffold protein nu3 (M309I and T310A), specifically mapping to the C-terminal region of gpC (*18*) (Fig. 5C). These findings are consistent with previous work identifying phage prohead proteases as key determinants of type I CBASS activation (*11*), suggesting that phage protease may serve as a conserved trigger for bacterial CD-NTase signaling.

**Fig. 5.**
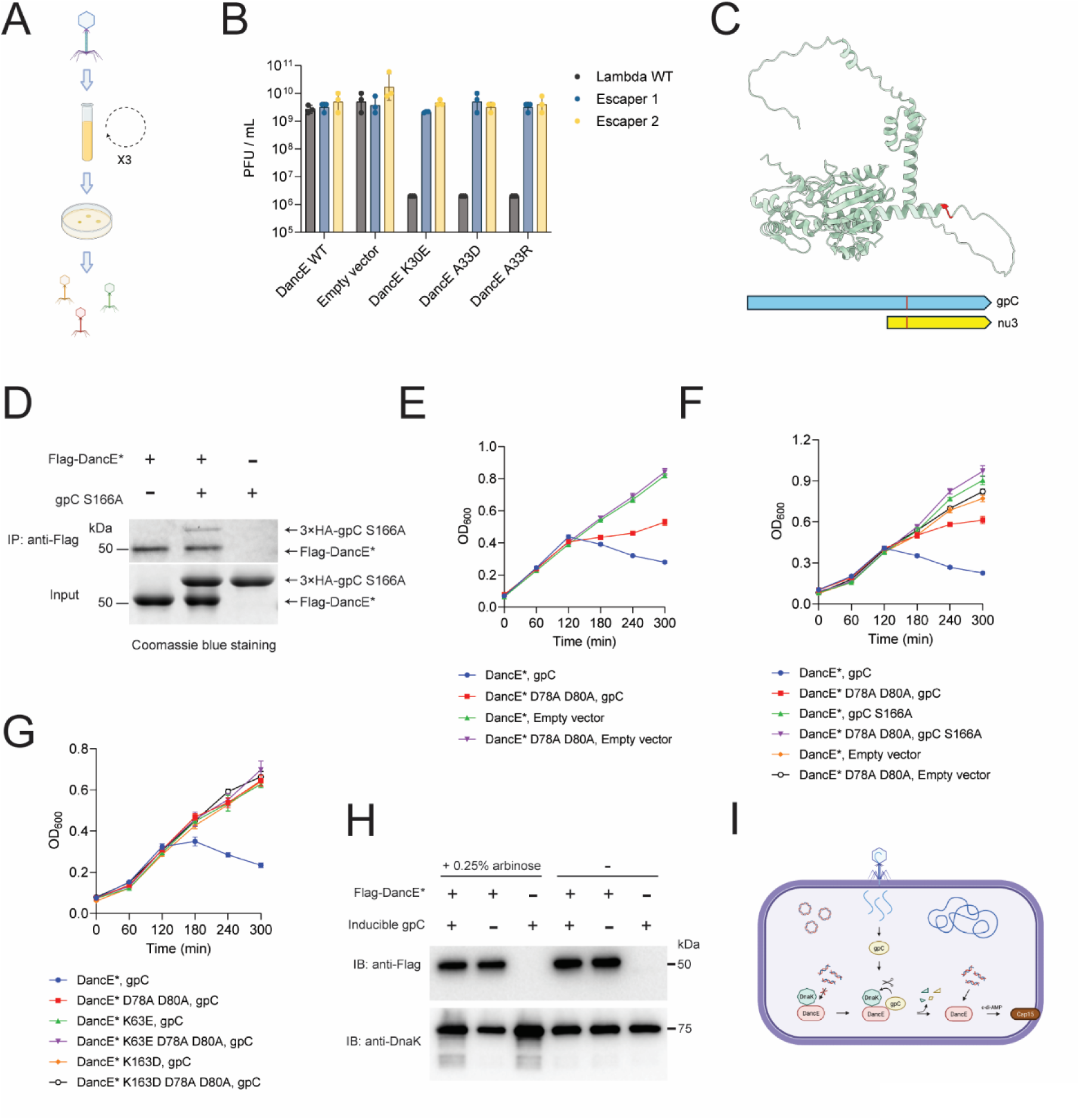
Lambda phage protease gpC triggers DancE activation. **(A)** Forward genetic screen for lambda phage escapers that evade DancE A33R mutant. Created with BioRender. (**B**) Efficiency of plating (EOP) of wild-type and escaper lambda phages on *E. coli* MG1655 expressing wild type operon, empty vector (no operon), or K30E, A33D and A33R mutants of DancE CBASS. Data are mean ± s.d. from three biological replicates. (**C**) Predicted structure of lambda phage gpC generated by AlphaFold 3 (pTM = 0.61) and visualized in ChimeraX. Escaper mutation sites are highlighted in red. (**D**) Immunoprecipitation analysis of DancE-gpC interaction. Purified Flag–DancE A33R (Flag-DancE*) and 3×HA-gpC S166A proteins were incubated with anti-Flag M2 magnetic beads (Sigma) for immunoprecipitation analysis. Analyzed by SDS–PAGE and Coomassie blue staining. (**E**)-(**G**), Reconstituted CBASS activation monitored by cellular growth curves. (**E**) Effects of wild type gpC or empty vector on the growth of cells expressing DancE A33R (DancE*) or the catalytically inactive DancE* D78A D80A mutant. (**F**) Effects of wild type gpC, the catalytically inactive gpC S166A mutant, or empty vector on the growth of cells expressing DancE* or DancE* D78A D80A mutant. (**G**) Effects of wild-type gpC or empty vector on the growth of cells expressing DancE*, DNA-binding mutants of DancE* (K63E or K163D), or the corresponding catalytically inactive D78A D80A mutants. Data are mean ± s.d. from three biological replicates. (**H**) Western blot analysis of Flag-DancE* and DnaK following 4 h of gpC induction with 0.25% arabinose. DancE A33R carrying an N-terminal Flag tag were expressed under the control of the native promoter, Cap15 was deleted in the operon. Anti-Flag antibody (Sigma) and anti-DnaK antibody (Abcam) were used for immunoblotting. (**I**) Schematic model of the proposed mechanism regulating DancE activation. Under basal conditions, DnaK suppresses activation of DancE by cellular DNA. During phage infection, phage-derived factors, such as lambda gpC, relieve DnaK-mediated repression, enabling DNA-dependent activation of DancE and the induction of abortive infection. Created with BioRender.

Because wild-type gpC undergoes autoproteolysis (*18*), we were unable to purify the wild-type gpC protein. Therefore, we used the catalytically inactive mutant gpC S166A (*18*) to examine its interaction with DancE. Immunoprecipitation assay showed that both DancE A33R and wild type proteins bind gpC S166A protein (Fig. 5D and fig. S8A). To directly test whether lambda gpC alone is sufficient to activate DancE, we employed a two-plasmid system in *E. coli*, expressing the gain-of-function DancE A33R mutant under its native promoter and gpC under an arabinose-inducible promoter. Induction of gpC expression was sufficient to trigger CBASS-mediated cellular toxicity (Fig. 5E and fig. S8B), demonstrating that gpC acts as a potent activator of DancE gain-of-function mutant *in vivo*. Importantly, the S166A mutation that abrogates the protease activity of gpC rendered the protein incapable of activating DancE, indicating the protease activity of gpC is required for DancE activation *in vivo* (Fig. 5F). We found that the gpC escaper mutants retained the ability to activate DancE A33R. Despite their escape phenotype, both gpC mutants remained capable of inducing CD-NTase activation (fig. S8C), suggesting the escaper mutations may have other effects on the phage, consistent with previous observations (*11*). Co-expression of the scaffold protein nu3 alone failed to activate DancE (fig. S8D), further supporting a specific role for gpC protease function. In contrast to DancE A33R variant, wild-type DancE was not activated by the lambda gpC, indicating that the lambda gpC specifically activates DancE A33R through a mechanism that requires further investigation (fig. S8E).

To determine whether gpC-mediated activation of DancE requires DNA binding, we introduced mutations disrupting the DNA-binding sites (K63E and K163D) of the DancE A33R mutant and co-expressed these variants with gpC. Disruption of DNA binding completely abolished gpC-induced activation of DancE (Fig. 5G), demonstrating that DNA binding by DancE is essential for its activation by the gpC protease activity.

We observed DnaK degradation products following gpC induction (Fig. 5H), whereas no DancE cleavage was detected despite the direct interaction between gpC and DancE (Fig. 5D). These results suggest that gpC binds DancE without cleaving it while selectively degrading DnaK. Given that gpC itself undergoes autoproteolysis (*18*), this process may subsequently release DancE to bind DNA. The increased DnaK expression upon gpC co-expression likely reflects a compensatory stress response induced by gpC-mediated proteostasis stress (*19*). Based on the results, we propose a model in which phage infection delivers the gpC protease that disrupts the interaction between DancE and host repressors such as DnaK, thereby allowing DancE to sense DNA and initiate an abortive infection response that limits phage propagation (Fig. 5I).

## Discussion

In this study, we identified a DNA-dependent activation mechanism for DancE from *Escherichia albertii* and demonstrate that this activation is repressed by the bacterial chaperon DnaK *in vitro*. We further show that mutation at the DancE dimer interface confers gain-of-function antiviral activity against specific bacteriophages, especially lambda phage, and that lambda phage prohead protease gpC activates DancE through a mechanism that depends on the protease activity of gpC. Together, our findings support a model in which DancE activation is centered on DNA sensing, a strategy that is evolutionarily conserved with mammalian cGAS-mediated innate immune signaling.

In mammalian cells, cGAS is primarily localized to the cytosol, while self-DNA is normally confined within the nucleus and mitochondria by membrane barriers to prevent aberrant cGAS activation under homeostatic conditions (*16, 20*). Upon viral infection or some cellular damage that delivers DNA into the cytoplasm, cGAS becomes activated, catalyzing the synthesis of cGAMP, which in turn activates the downstream adaptor STING to initiate antiviral and inflammatory responses (*16*).

Remarkably, despite the absence of membrane-bound organelles in bacteria, we find that DancE is similarly activated by DNA. This presents an apparent paradox: in bacterial cells, chromosomal and plasmid DNA are constitutively present in the cytosol and would be expected to continuously stimulate a DNA-sensing enzyme. Our finding that DnaK represses DancE by preventing its DNA binding provides a mechanism of DancE repression in the absence of infection. Following lambda phage infection, the phage prohead protease gpC activates DancE A33R through a mechanism that depends on the gpC protease activity. While this observation is consistent with our observation that lambda phage specifically activates DancE A33R, it is not clear how wild-type DancE is activated by other phages such as the T2 phage. Exploration of this question is hampered by our observation that the prohead protease from T2 phage is toxic to *E. coli* even in the absence of DancE.

We have attempted to determine if DnaK is the dominant repressor of DancE *in vivo* by using an inducible degradation system to deplete DnaK in *E. coli*. However, we found that depletion of DnaK using this method did not cause activation of DancE, suggesting other repressors of DancE may exist in cells. Although we could detect partial DnaK degradation in cells that express the gpC protease, we have not obtained evidence that this degradation causes DancE activation. It is possible that the gpC protease can target other repressors of DancE in addition to DnaK. Further research is needed to fully delineate the mechanism of DancE activation by phage infection. Nevertheless, our finding that the DancE can be activated by DNA and repressed by DnaK, and that the protease activity of gpC is required for DancE activation *in vivo* supports a guard hypothesis in which the immune sensor DancE is regulated by guard proteins such as DnaK that blocks its sensing of DNA. In response to phage infection, DancE indirectly “senses” the phage protease activity that de-represses the guard, enabling DancE activation by DNA. Our data show that DNA sensing by cGAS-like enzymes is evolutionarily conserved, but the regulatory mechanisms of cGAS activation are diverse and sophisticated across kingdoms of life.

## ACKNOWLEDGEMENTS

Use of the Stanford Synchrotron Radiation Lightsource, SLAC National Accelerator Laboratory, is supported by the U.S. Department of Energy, Office of Science, Office of Basic Energy Sciences under Contract No. DE-AC02-76SF00515. The SSRL Structural Molecular Biology Program is supported by the DOE Office of Biological and Environmental Research, and by the National Institutes of Health, National Institute of General Medical Sciences (P30GM133894). The contents of this publication are solely the responsibility of the authors and do not necessarily represent the official views of NIGMS or NIH. We thank the Structural Biology Laboratory at UT Southwestern Medical Center for support with X-ray crystallographic studies. We thank the Macromolecular Biophysics Resource at UT Southwestern Medical Center for support with mass photometry experiments. The mass photometer was supported by NIH award S10OD030312-01.

## Author contributions

Experiments were conceived and designed by Y.L., M.H., and Z.J.C. Phage and bacterial experiments were performed by Y.L. Protein purification and *in vitro* enzymatic assays were performed by M.H. Mass spectrometry experiments were performed Y.L. *In vitro* biochemical assays were performed by Y.L. and M.H. X-ray data collection/analyses were performed by M.H., J.M.J. and D.R.T. Z.J.C. supervised the study. Z.J.C., Y.L. and M.H. wrote and revised the manuscript with input from all authors. All authors contributed to editing the manuscript and supporting the conclusions.

## Data and materials availability

The structures of apo DancE and THFA-bound DancE have been deposited in the RCSB Protein Data Bank under accession codes 37ZT and 37ZU, respectively. The *E. coli* strain K12 proteome database that was used in this study can be accessed at Uniprot (UP000000625). Sequence Read Archive (SRA) under BioProject PRJNA1457026. IMG accessions for the CD-NTases and CBASS operon used in this study are found in the Methods. Materials, including strains and plasmids, are available upon reasonable request. Source data are provided with this paper.

## Supplementary Materials

### Materials and Methods

#### CD-NTase library in HEK 293T cells

Sixteen CD-NTases were selected based on previous reports demonstrating *in vitro* activity (*1–4*) (table S1). The corresponding coding sequences were commercially synthesized and cloned into the pcDNA3.1 vector. HEK 293T cells at approximately 60% confluence in 6- well plates were transfected with 2 μg of plasmid per well using polyethylenimine (PEI). Cell phenotypes were observed 48 h after transfection. Cells were then harvested, and small molecules were extracted using 80% of methanol, then purified by Hypersep Spe Nh2 column (Thermo) and analyzed by LC-MS. The spectrum data were checked and analyzed by Xcalibur Qual Browser (Thermo), and pie figure was generated by GraphPad Prism according to peak area of different products.

#### Cloning DancE CBASS into *E. coli*

The cloning strategy was adapted from a previous study (*5*). Briefly, the DancE CBASS operon, together with approximately 450 bp of flanking sequence on each side containing the putative native promoter and terminator, was commercially synthesized based on the following sequence: *Escherichia albertii* (IMG: Ga0062141_1142, 1-2830). The synthesized fragment was assembled into an empty vector containing the p15A origin of replication and a chloramphenicol resistance cassette using Gibson assembly (*6*). Site-mutant plasmids were also generated by Gibson assembly. A Flag tag for expression level examination was added to the N-terminal of the DancE coding sequence. DH5α cells were used for plasmid amplification, and the plasmids were verified by whole plasmid sequencing. The plasmids were transformed into *E. coli* MG1655 or BW25113 for indicated experiments.

#### Protein alignments

Twenty randomly selected AGS-C domain-containing type I CD-NTase sequences from a previously reported study (*4*) were aligned with DancE. The sequence logo of conserved sites were displayed using WebLogo (*7*). DancE was aligned with DancE homologs (IMG gene ID 2746149570, 2611579298 and 2526020757) using Clustal W (*8*), and the alignment was visualized by ESPript (*9*).

#### Recombinant protein expression and purification

Full-length wild-type DancE and the indicated mutants were used for *in vitro* assays, and wild-type DancE was used for crystallographic studies. *Escherichia coli* DnaK, human cGAS and *Escherichia coli* DncV were used for *in vitro* assays. All proteins were expressed in *Escherichia coli* BL21(DE3) pLysS cells using a pET-SUMO expression vector. Starter cultures were grown in terrific broth (TB; RPI) at 37 °C to an OD_600_ of ∼0.6, followed by induction with 1 mM IPTG and overnight expression at 18 °C.

For apo-DancE and human cGAS purification, cells were harvested by centrifugation and resuspended in lysis buffer containing 20 mM Tris-HCl (pH 8.0), 500 mM NaCl, 2% (v/v) glycerol, 20 mM imidazole, and 0.2 mM phenylmethylsulphonyl fluoride (PMSF). Cells were lysed by sonication, and the lysate was clarified by centrifugation. The supernatant was incubated with Ni-NTA agarose resin (Qiagen) at 4 °C for 2 h with gentle rotation.

The resin was washed extensively with lysis buffer, and bound proteins were eluted with elution buffer containing 20 mM Tris-HCl (pH 8.0), 500 mM NaCl, 2% glycerol, and 300 mM imidazole. The N-terminal SUMO tag was removed by incubation with ULP1 protease (1:100, w/w) at 4 °C overnight in the presence of 1 mM DTT, accompanied by dialysis into buffer containing 20 mM Tris-HCl (pH 8.0), 500 mM NaCl, 2% glycerol, and 1 mM DTT. The cleaved sample was passed through Ni-NTA resin again to remove the His_6_-SUMO tag and ULP1 protease. The untagged DancE protein was further purified by sequential ion-exchange chromatography using a HiTrap Heparin HP column (Cytiva) followed by a HiTrap Q HP column (Cytiva), both developed with a linear gradient of 0– 1 M NaCl in 25 mM Tris-HCl (pH 8.0). Final polishing was performed by size-exclusion chromatography on a Superdex 200 Increase 10/300 GL column (Cytiva) equilibrated in 10 mM Tris-HCl (pH 8.0) and 100 mM NaCl. For purification of DancE used to obtain the THFA-bound structure, both ion-exchange chromatography steps were omitted, and the protein was subjected directly to size-exclusion chromatography after the second Ni-NTA purification step. Purified protein was concentrated, flash-frozen in liquid nitrogen, and stored at −80 °C.

For DnaK and DncV purification, cells were resuspended in lysis buffer containing 25 mM Tris-HCl (pH 8.0), 150 mM NaCl, 20 mM imidazole, and 0.2 mM PMSF, lysed by sonication, and clarified by centrifugation. The supernatant was incubated with Ni-NTA resin at 4 °C for 2 h, washed with lysis buffer, and eluted with buffer containing 25 mM Tris-HCl (pH 8.0), 150 mM NaCl and 300 mM imidazole. The SUMO tag was cleaved with ULP1 protease (1:100, w/w) overnight at 4 °C in the presence of 1 mM DTT, with simultaneous dialysis into 25 mM Tris-HCl (pH 8.0), 30 mM NaCl, and 1 mM DTT. The sample was re-applied to Ni-NTA resin to remove the tag and protease. The untagged DnaK protein was further purified by anion-exchange chromatography on a HiTrap Q HP column (Cytiva) using a 0–1 M NaCl gradient in 25 mM Tris-HCl (pH 8.0), followed by size-exclusion chromatography on a Superdex 200 Increase 10/300 GL column (Cytiva) equilibrated in 10 mM Tris-HCl (pH 8.0) and 100 mM NaCl. The purified protein was concentrated, flash-frozen, and stored at −80 °C.

#### X-ray crystallography and structure determination

Crystals of full-length wild-type apo-DancE were obtained using the hanging-drop vapor diffusion method at 18 °C by mixing 1 μl of protein solution (5 mg ml−1) with 1 μl of reservoir solution. Diffraction-quality crystals grew in a reservoir containing 1.0 M succinic acid (pH 7.0), 0.1 M HEPES (pH 7.0), and 1.6% (w/v) polyethylene glycol monomethyl ether 2000 (PEG MME 2000). For THFA-bound DancE, 1 μl of protein solution was mixed with 1 μl of reservoir solution containing 1.2 M ammonium sulfate, 0.1 M Tris-HCl (pH 8.0), and 10% (v/v) glycerol. DancE crystals were cryoprotected by supplementing the reservoir solution with 15% (v/v) ethylene glycol and flash-cooled in liquid nitrogen prior to data collection. X-ray diffraction data were collected at beamline BL12-2 at the Stanford Synchrotron Radiation Lightsource (SSRL), SLAC National Accelerator Laboratory (Menlo Park, CA, USA). Diffraction data for apo- and THFA- bound DancE crystals were indexed, integrated, and scaled using HKL-3000. The structure was solved by molecular replacement using PHASER with an AlphaFold-predicted model of DancE as the search model (*10–12*). Model building and refinement were performed through iterative cycles using Coot and PHENIX (*13, 14*). Data collection and refinement statistics are summarized in table S2. Structural figures were prepared using UCSF ChimeraX (v1.9).

#### Mass photometry

Mass photometry measurements were performed using a Refeyn TwoMP mass photometer. For each measurement, 16.2 µl of protein buffer (10mM Tris, PH 8.0, 100mM NaCl) was applied to a clean coverslip, and the instrument was focused using the droplet-dilution mode. DancE was then added to the droplet and mixed immediately to a final volume of 18 µl and a final protein concentration of 100 nM. Movies were recorded for 60 s using AcquireMP software (version 2026.1.0.0; Refeyn). Molecular mass distributions were obtained from the recorded movies and analyzed using DiscoverMP software (version 2026.1.1.0; Refeyn).

#### Phage propagation

Phages used in this study were propagated in *E. coli* BW25113. Host cells were grown overnight in LB medium at 37 °C with shaking (220 rpm). The overnight culture was diluted 1:100 into fresh LB medium supplemented with 5 mM MgCl_2_, 5 mM CaCl_2_ and 0.1 mM MnCl_2_. Phages were inoculated when the OD_600_ of *E. coli* culture reached 0.2. The culture was continueduntil the cells completely lysed. Several drops of chloroform were added to the media to facilitate cell lysis, and the debris was then removed by centrifugation at 4,000 rpm for 10 min. The supernatant containing the phages was transferred to a new tube, supplemented with a few additional drops of chloroform, and stored at 4 °C.

#### Efficiency of plating (EOP) assay

Phage titers, the efficiency of plating (EOP), and library screening were determined using small-drop plaque assays. Briefly, 120 μl of overnight bacterial culture was mixed with 5 ml of molten top agar (0.5% agar in LB, supplemented with 5 mM MgCl_2_, 5mM CaCl_2_ and 0.1 mM MnCl_2_, kept at 50 °C), which was then quickly poured onto a Petri dish. The plates were kept at room temperature for 30 minutes before plaque assays. Tenfold serial dilutions of phage stocks were prepared in LB, and 5 μl of each dilution was spotted onto the bacterial lawn. Plates were incubated overnight at 37°C. Plaque-forming units (PFU) were quantified by counting individual plaques at the highest dilution that produced visible lysis. When plaques were too small to distinguish individually, the spot was counted as 10 PFU. Experiments were performed in triplicate to demonstrate reproducibility. Data analysis was performed by GraphPad Prism.

#### Immunoblotting

Bacterial cultures were harvested by centrifugation and resuspended in 1× LDS sample buffer supplemented with 5% β-mercaptoethanol. Samples were boiled for 10 min and centrifuged at 13,000 rpm for 10 min to remove debris. Proteins were separated by SDS– PAGE using 4–20% Tris–glycine gels (Novex) and transferred onto PVDF membranes (Bio-Rad, 1704157). Membranes were blocked with 5% non-fat milk in TBST (20 mM Tris-HCl, pH 7.5, 150 mM NaCl, 0.1% Tween-20). For detection of Flag–DancE, membranes were incubated with an anti-Flag antibody (1:2,000; Sigma), followed by an HRP-conjugated anti-mouse IgG secondary antibody (1:2,000; Cell Signaling Technology). For detection of DnaK, membranes were incubated with an anti-DnaK antibody (1:2,000; Abcam), followed by the same HRP-conjugated secondary antibody. Chemiluminescent signals were acquired using a ChemiDoc imaging system (Bio-Rad).

#### DancE Enzymatic Assay

For *in vitro* enzymatic assays, recombinant DancE, DncV or cGAS proteins were incubated with or without the indicated DNA (plasmid used for enzymatic assay is pcDNA3.1) or DnaK in reaction buffer (10mM Tris-HCl pH 7.5, 25mM KCl, 20mM MgCl_2_, 1mM MnCl_2_, 1mM DTT, 2mM each of rNTP) at 37 °C for the indicated times. Reactions were terminated by adding four volumes of methanol, followed by centrifugation to remove precipitated material. The supernatants were evaporated to dryness using a SpeedVac, and the dried extracts were resuspended in water for LC–MS analysis. C-di-AMP concentrations were determined using a serial dilution of c-di-AMP standards. Assays were performed in triplicate to ensure reproducibility.

A QExactive HF-X quadrupole-orbitrap hybrid mass spectrometer (Thermo) linked to Vanquish UPLC (Thermo) was used to detect small molecules. An amide column (ACQUITY UPLC BEH Amide 1.7µm 1.0×150mm, Waters) was used for separation with 0.1% formic acid in acetonitrile as mobile phase A and 0.1% formic acid as mobile phase B. The gradient was set as follows with flow rate being set at 80 µl/min: 5% B at 0-2 min, 60% B at 21.8 min, 60% B at 21.8-24.8 min, 5% B at 24.9 min, 5% B at 24.9-33 min. Analytes were sprayed with electrospray Ionization (ESI) source at a spray voltage of 3.5 kV and a capillary temperature of 250 °C. Positive model data-dependent acquisition was used for data collection with full-scan MS was acquired at an m/z range of 300–1,200, resolution of 120,000 and automatic gain control target of 3 × 10^6^ ions. The top 5 most intense ions were selected for high energy collisional dissociation fragmentation, and the resolution was set at 30,000. Automatic gain control target was set at 1 × 10^5^. Stepped collision energies were set at 25, 30, 35 eV. Dynamic exclusion was set at 10 s. The spectrum data were checked and analyzed by Xcalibur Qual Browser (Thermo). Data analysis was performed by GraphPad Prism.

#### 5-hmc DNA fragment preparation

A 380-bp fragment containing the CMV enhancer from pcDNA3.1 was amplified by PCR using either standard deoxyribonucleotides (dNTPs) or dNTPs in which dCTP was replaced with 5-hydroxymethyl-dCTP (5-hm-dCTP). The resulting PCR products were purified and used for enzymatic assays.

#### Electrophoretic mobility shift assay (EMSA)

The indicated proteins were incubated with 5 ng/µl linearized pcDNA3.1 plasmid in reaction buffer containing 10 mM Tris-HCl (pH 7.5), 75 mM KCl, and 1 mM MgCl_2_ at 37 °C for 20 min. The reaction mixtures were then resolved on a 1.5% agarose gel containing 1 µg/ml ethidium bromide in 1× TBE buffer. Gels were imaged using a ChemiDoc Imaging System (Bio-Rad).

#### Isolation and identification of cyclic dinucleotides produced by DancE in vivo

For *in vivo* products detection, *E. coli* BW25113 cells were transformed with plasmids encoding the DancE A33R mutant under an arabinose-inducible promoter and N-terminally Flag-tagged truncated Cap15 (residues 69–195) under the constitutive J23119 (BBa_J23119) promoter (*15*). Cultures were grown overnight, diluted 1:100 into fresh medium, and grown to an OD_600_ of 0.4. Cells were then harvested and lysed by sonication. Flag-tagged Cap15 was immunoprecipitated using anti-Flag M2 magnetic beads (Sigma) to isolate Cap15-bound small molecules. The bound small molecules were extracted with 80% methanol and analyzed by LC–MS. Assays were performed in triplicate to ensure reproducibility. Data analysis was performed by GraphPad Prism. Pie figure was generated according to peak area of different products.

#### Protein identification by mass spectrometry

Protein band containing DancE inhibiting factor was excised and de-stained with de- staining buffer (25 mM NH_4_HCO_3_, 50% acetonitrile), followed by reduction using 5mM dithiothreitol and alkylation using 50 mM iodoacetamide. The gel was then digested with 12.5 ng/μl trypsin (Promega) at 37 °C overnight. The resulting peptides were extracted with 1% formic acid and purified with Sep-Pak C18 cartridge (Waters) before analyzed by LC-MS.

An in-house packed C18 column in silica capillary emitters (resin: 100 Å, 3 μm, MICHROM Bioresources; column: 100 μm internal diameter, 100 mm resin length) equipped on Vanquish UPLC (Thermo) was used to separate peptides. Flow rate was set at 600 nl/min at 0–13.5 min and 250 nl/min at 13.5–120 min with 0.1% formic acid as mobile phase A and 0.1% formic acid in acetonitrile as mobile phase B. The gradient was set as follows: 2% B at 0–15 min, 30% B at 81 min, 35% B at 85 min, 40% B at 87 min, 60% B at 95 min, 80% B at 96–107 min and 2% B at 108–120 min.

A nano-electrospray Ionization (nESI) source equipped on QExactive HF-X quadrupole-orbitrap hybrid mass spectrometer (Thermo) was used for peptide ionization. The spray voltage was set at 1.5 kV and capillary temperature was set at 250 °C. Positive model data-dependent acquisition was used for data collection with full-scan MS was acquired at an m/z range of 300–1,650, resolution of 60,000 and automatic gain control target of 3 × 10^6^ ions. The top 15 most intense ions were selected for high energy collisional dissociation fragmentation, and the resolution was set at 15,000. Automatic gain control target was set at 1 × 10^5^. Collision energies were set at 30 eV. Dynamic exclusion was set at 30 s.

SEQUEST algorithm in Proteome Discoverer (Thermo) was used for peptide-spectrum matching with the *E. coli* K12 proteome database (Uniprot, UP000000625). Carbamidomethylation on cysteines was set as static modification. Methionine oxidation was set as variable modification. Precursor mass error was set as 10 ppm and fragment mass error was set at 0.05 Da. Maximum mis-cleavage was set at 2. Peptide false discovery rate was set at 1%. A complete list of protein identified for DnaK identification is available in table S3.

#### Immunoprecipitation assay

For the DancE–DnaK immunoprecipitation assay, purified Flag-DancE (4 µM) and DnaK (1.6 µM) were incubated together. For the DancE–gpC immunoprecipitation assay, purified Flag-DancE WT or A33R (4 µM) was incubated with HA-tagged gpC S166A (4 µM). Proteins were incubated with anti-Flag M2 magnetic beads (Sigma) in buffer containing 25 mM Tris-HCl (pH 8.0), 300 mM NaCl, 0.1% Tween-20, and 5% glycerol at 4 °C for 2 h with gentle rotation. The beads were subsequently washed three times with the same buffer, and bound proteins were eluted with 3×Flag peptide. Eluted proteins were resolved by SDS–PAGE using 4–20% Tris–glycine gels (Novex) and visualized using InstantBlue Coomassie Protein Stain (Abcam, ab119211).

#### AlphaFold 3 structure prediction

For prediction of the DancE–DNA complex, two copies of full-length DancE and two copies of a 15-bp dsDNA (TACAGATCTACTAGT) were used. For DancE homolog– DNA complexes, predictions were performed using two copies of full-length CD-NTase from *Escherichia coli* KCJK1734 (IMG gene ID 2746149570) with two copies of the 15- bp dsDNA, two copies of full-length CD-NTase from *Pectobacterium parmentieri* RNS08.42.1A (IMG gene ID 2611579298) with two copies of the 15-bp dsDNA, or two copies of full-length CD-NTase from *Alteromonas* sp. ALT199 (IMG gene ID 2526020757) with two copies of a 20-bp dsDNA (TACAGATCTACTAGTGATCT). For prediction of the Cap15–c-di-AMP complex, one molecule of full-length wild-type Cap15 from the DancE CBASS system and one molecule of bis-(3’,5’)-cyclic-dimeric-adenosine monophosphate (3’3’-c-di-AMP) were used. For prediction of the λ phage gpC structure, one copy of full-length gpC was used. All structures were predicted using the AlphaFold 3 web server (alphafoldserver.com)(*12*).

#### Lambda phage mutagenesis and screening

*E. coli* MG1655 harboring the DancE A33R CBASS operon was grown overnight and diluted 1:100 into fresh LB medium supplemented with 5 mM MgCl_2_, 5 mM CaCl_2_, and 0.1 mM MnCl_2_. Wild type λ phage was added when the culture reached an OD_600_ of 0.2. After 24 h of incubation, the resulting phage lysate was used to infect a fresh culture of *E. coli* MG1655 harboring DancE A33R CBASS. This procedure was repeated for a total of three rounds of phage propagation. Following the third round, phage lysates were subjected to plaque assays using *E. coli* MG1655 harboring DancE A33R CBASS as the host. Individual large plaques, representing candidate phage escapers, were isolated and propagated. The escape phenotype was subsequently confirmed by plaque assay.

To determine the genotypes of the phage escapers, phage genomic DNA was extracted using a phage DNA isolation kit (Norgen Biotek, NC0690454). Whole-genome sequencing of the escapers and wild type λ phage was performed on an Illumina NovaSeq X Plus platform with 150-bp paired-end reads. Sequencing reads were subjected to quality control and subsequently assembled using SeqMan NGen (DNASTAR).

#### Bacterial growth curve

*E. coli* BW25113 cells transformed with the indicated protein-expression plasmids were grown overnight at 37 °C and diluted 1:20 into fresh LB medium. This time point was designated as 0 min, and protein expression was induced by the addition of 0.25% (w/v) arabinose. OD_600_ was measured every hour using a CLARIOstar plate reader (BMG Labtech). Data analysis was performed by GraphPad Prism.

**Fig. S1.**
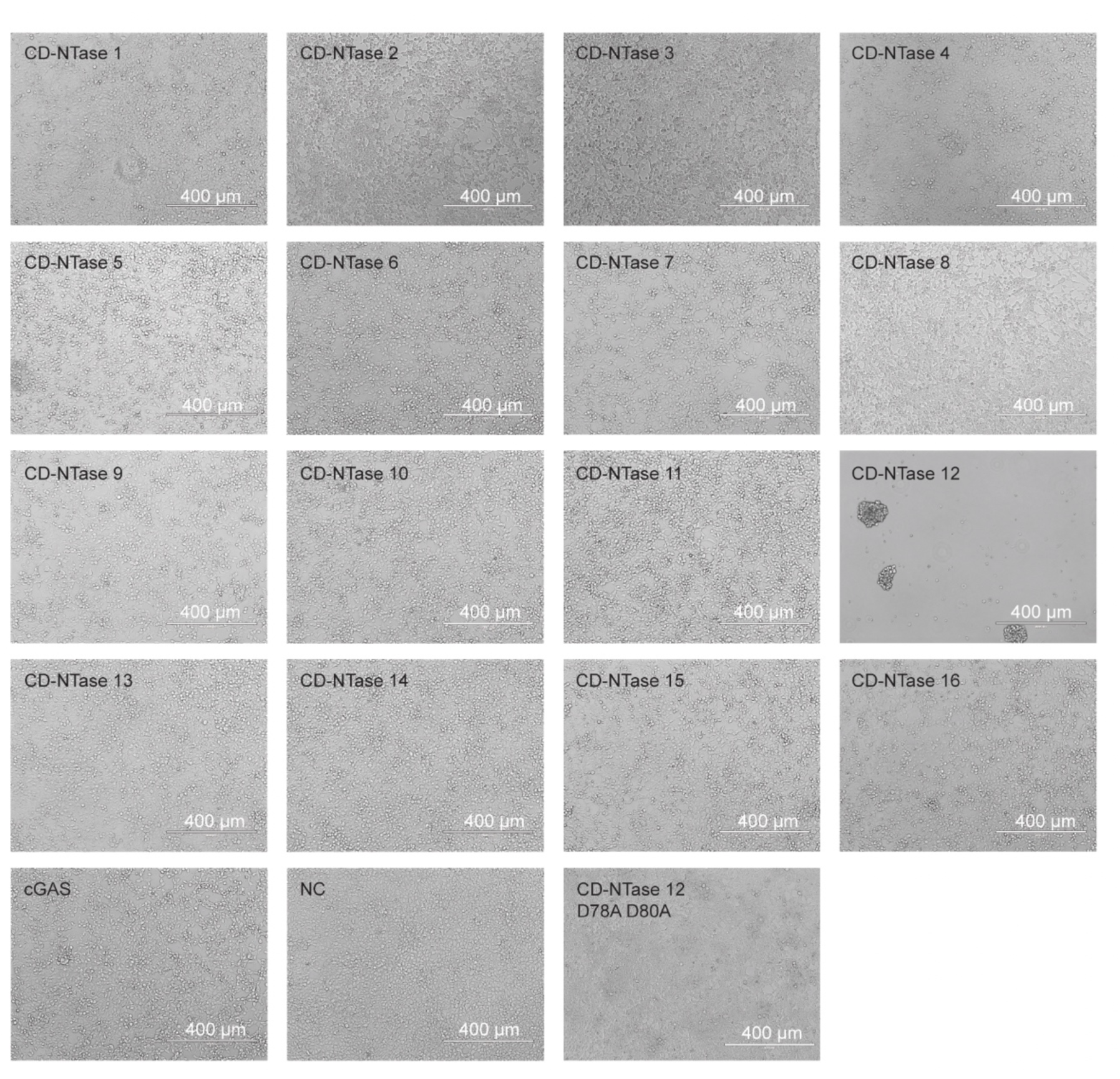
HEK 293T cell morphology following transfection with different CD-NTases. HEK 293T cells in 6-well plates were transfected with 2 μg of plasmid per well using polyethylenimine (PEI). Cell phenotypes were observed at 48 h after transfection.

**Fig. S2.**
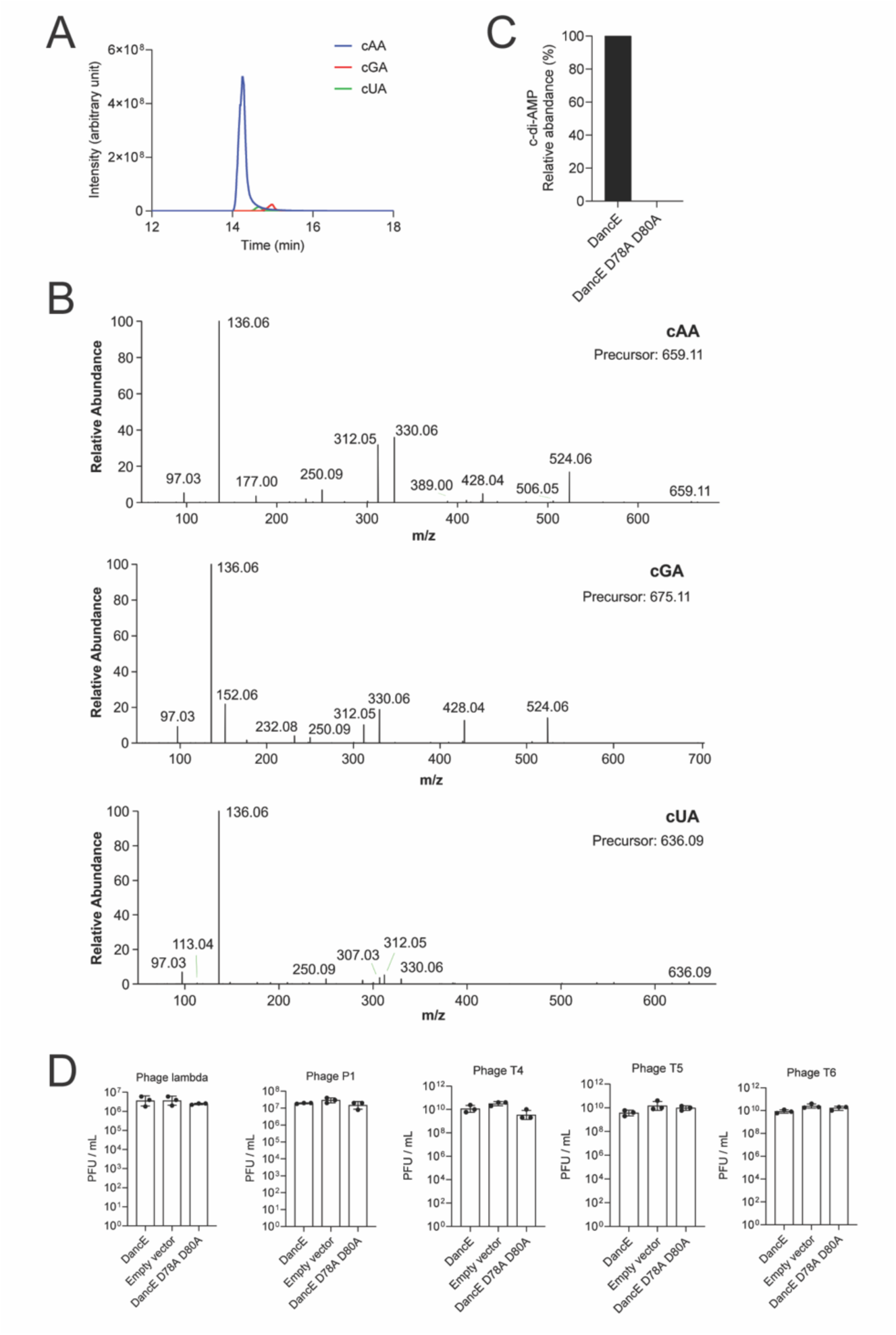
Identification of the products of DancE in HEK 293T cells and antiphage function of DancE. (**A**) Chromatographic comparison of the DancE products in HEK 293T cells. Separation was performed using an amide column (ACQUITY UPLC BEH Amide, 1.7 µm, 1.0 × 150 mm; Waters). cAA, c-di-AMP; cGA, c-GMP-AMP; cUA, c-UMP-AMP. (**B**) MS/MS fragmentation spectra of the DancE products in HEK 293T cells. (**C**) DancE catalytic dead mutant (D78A D80A) didn’t show enzymatic activity in HEK 293T cells. The data are representative of at least three independent experiments. (**D**) Efficiency of plating (EOP) of lambda, P1, T4, T5 and T6 phages on *E. coli* MG1655 expressing wild-type operon, empty vector (no operon), or D78A D80A mutant (DancE catalytic dead) of DancE CBASS. Data are mean ± s.d. from three biological replicates.

**Fig. S3.**
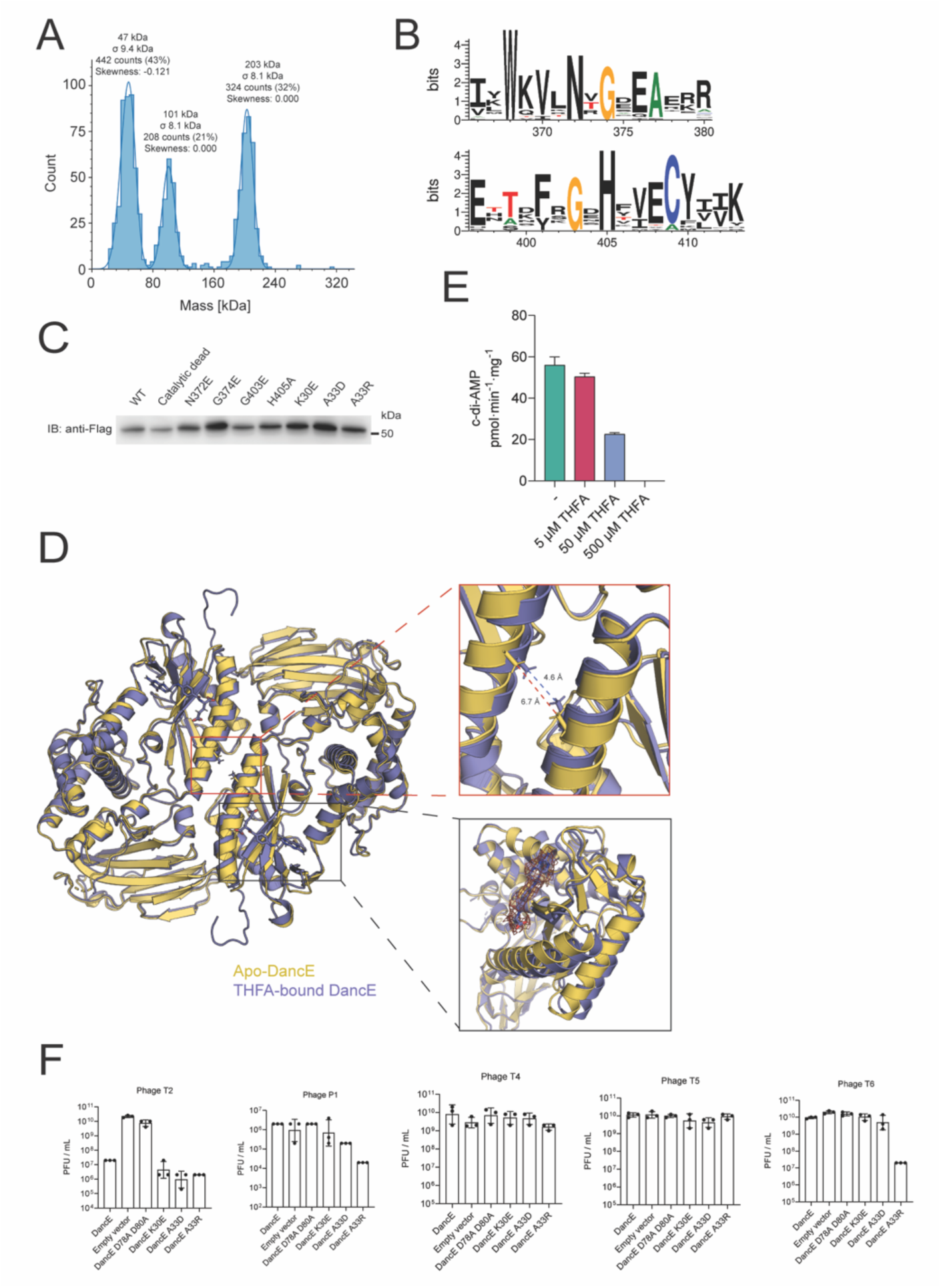
Structural basis of DancE architecture and its anti-phage function. (**A**) Mass photometry analysis of DancE. DancE (100 nM) showed three major populations centered at 47, 101, and 203 kDa, corresponding to monomeric, dimeric, and tetrameric forms, respectively. The data are representative of at least three independent experiments. (**B**) Sequence logo of aligned AGS_C domain regions containing conserved residues. (**C**) *In vivo* expression of DancE proteins. Wild-type DancE and the indicated DancE mutants carrying an N-terminal Flag tag were expressed under the control of the native promoter. Anti-Flag antibody (Sigma) was used for immunoblotting. (**D**) Structural alignment of THFA- bound DancE and apo-DancE crystal structures, RMSD = 1.044. Left, alignment of the dimer structures. Upper right, distances (Å) between the protomers in the THFA-bound DancE and apo-DancE dimers. Lower right, the α-helices of DancE-THFA and apo DancE are positioned slightly differently, with the electron density corresponding to THFA shown in the THFA-binding pocket. (**E**) Wild-type DancE protein (4 µM) was incubated with or without indicated concentration of THFA overnight (14 h). Data are mean ± s.d. from three independent replicates. (**F**) Efficiency of plating (EOP) of T2, P1, T4, T5 and T6 phages on *E. coli* MG1655 expressing wild type operon, empty vector (no operon), D78A D80A mutant (DancE catalytic dead), K30E, A33D, or A33R mutants of DancE CBASS. Data are mean ± s.d. from three biological replicates.

**Fig. S4.**
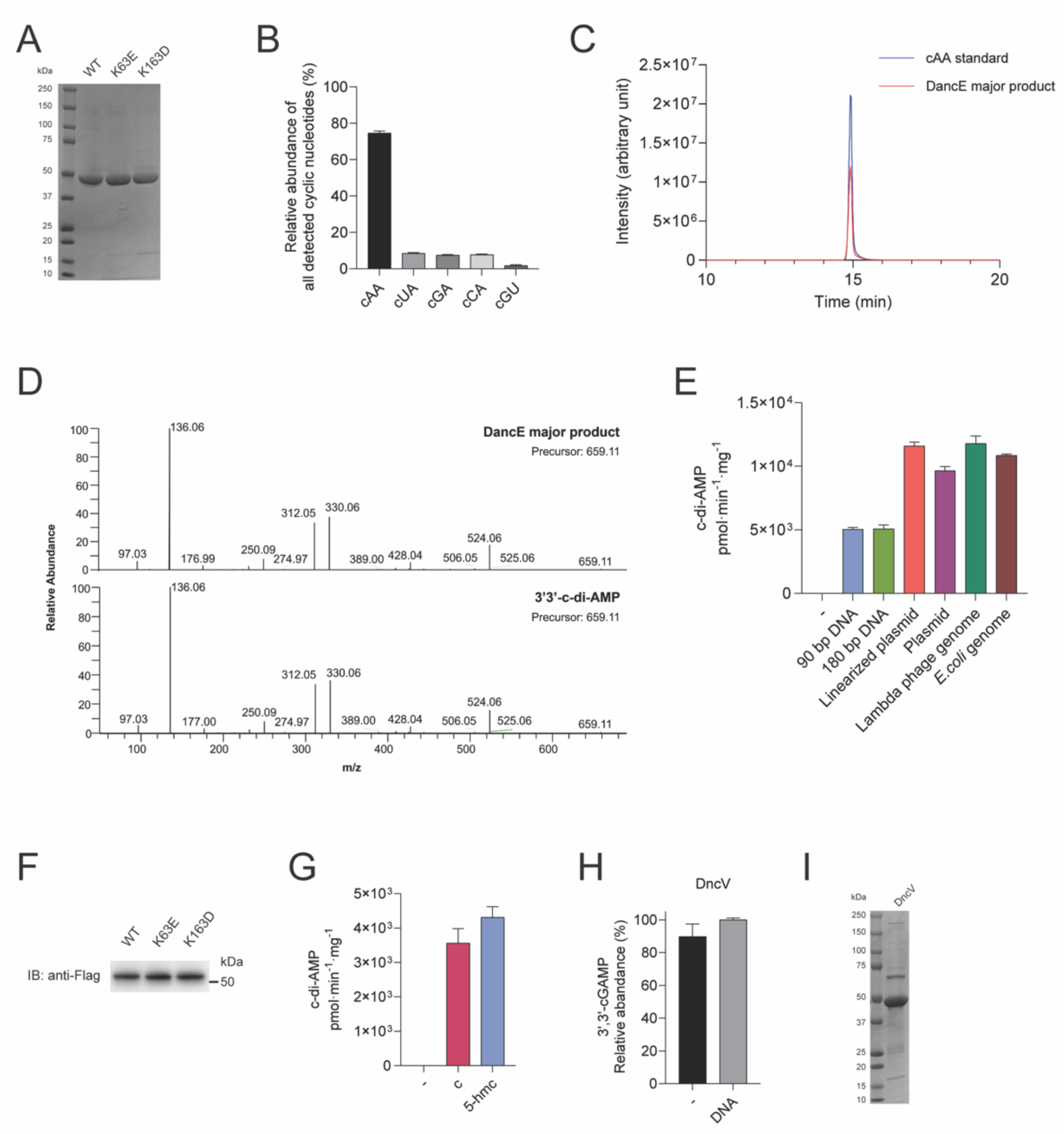
DNA activates DancE to trigger anti-phage signaling. (**A**) Purified recombinant wild-type and DNA-binding mutant DancE proteins analyzed by SDS–PAGE and Coomassie blue staining. (**B**)-(**D**), Characterization of DancE *in vitro* reaction products. Wild-type DancE protein (1 µM) was incubated with 45-bp dsDNA (275 ng/µl) for 2 h. (**B**) Quantification of DancE products showing c-di-AMP as the major product. cAA, c-di-AMP; cUA, c-UMP-AMP; cGA, c-GMP-AMP; cCA, c- CMP-AMP; cGU, cGMP-UMP. Data are mean ± s.d. from three independent replicates. (**C**) Chromatographic comparison of the major DancE reaction product and c-di-AMP standard (10 µM). Separation was performed using an amide column (ACQUITY UPLC BEH Amide, 1.7 µm, 1.0 × 150 mm; Waters). (**D**) MS/MS fragmentation spectra of the major DancE reaction product and the c-di-AMP standard. (**E**) Effects of DNA length on DancE activation. Wild type DancE protein (0.8 µM) was incubated with or without the indicated DNA (5 ng/µl) for 2 h. Data are mean ± s.d. from three independent replicates. (**F**) *In vivo* expression of DancE proteins. Wild-type DancE and the indicated DancE mutants carrying an N-terminal Flag tag were expressed under the control of the native promoter. Anti-Flag antibody (Sigma) was used for immunoblotting. (**G**) Wild-type DancE (0.8 µM) was incubated with or without 380-bp unmodified DNA (C) or 5-hydroxymethylcytosine (5-hmC)-modified DNA (1.4 ng/µl) for 2 h. Data are mean ± s.d. from three independent replicates. (**H**) Wild type DncV protein (0.8 µM) was incubated with or without plasmid (50 ng/µl) for 3 h. Data are mean ± s.d. from three independent replicates. (**I**) Purified recombinant DncV proteins analyzed by SDS–PAGE and Coomassie blue staining.

**Fig. S5.**
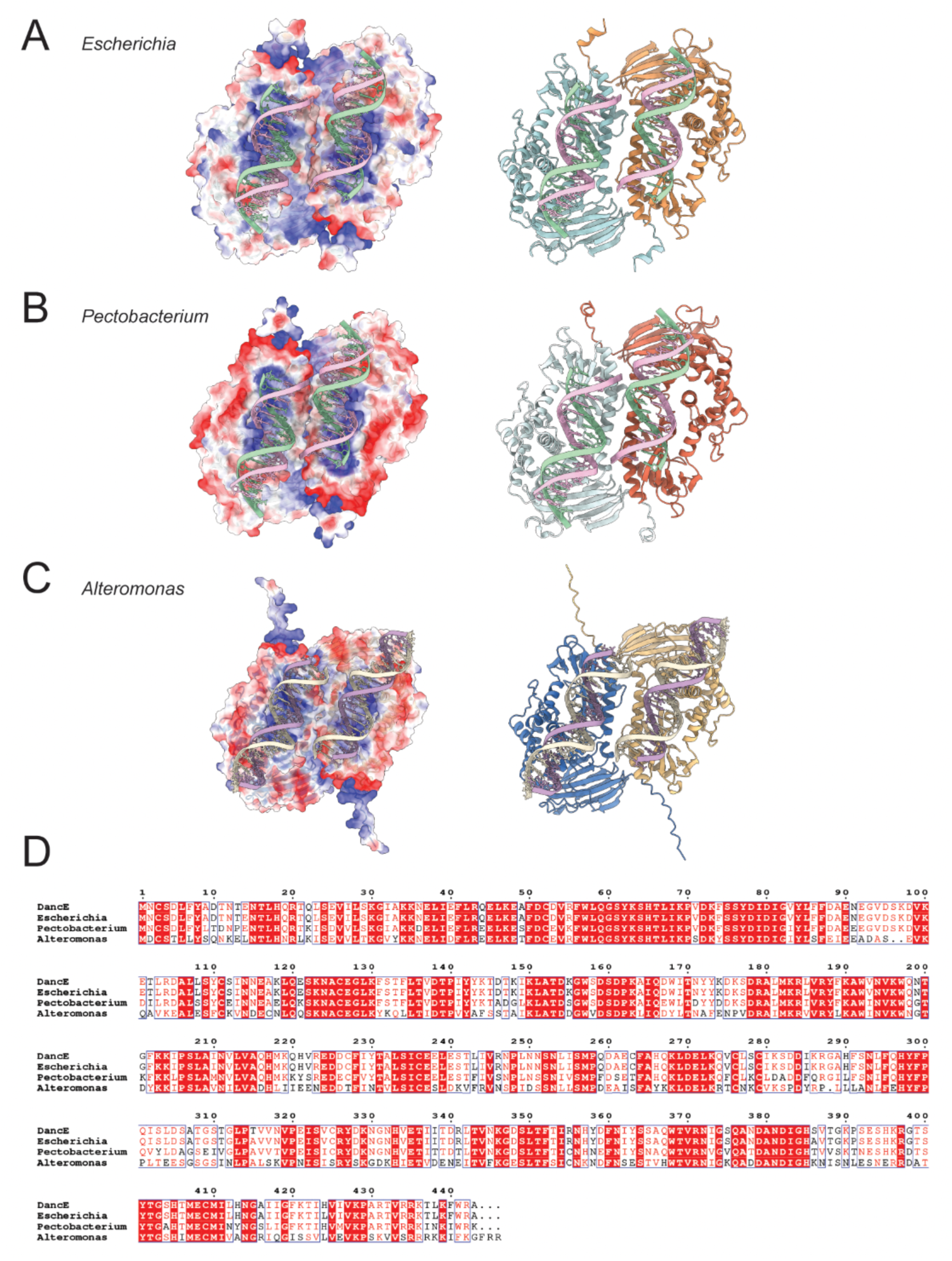
AlphaFold 3-predicted structures of DancE homolog-DNA complexes. (**A**)-(**C**), AlphaFold 3-predicted structures of DancE homolog–DNA complexes. (**A**) CD-NTase from *Escherichia coli KCJK1734* in complex with 15-bp DNA (ipTM = 0.82; pTM = 0.86). (**B**) CD-NTase from *Pectobacterium parmentieri RNS08.42.1A* in complex with 15-bp DNA (ipTM = 0.82; pTM = 0.85). (**C**) CD-NTase from *Alteromonas sp. ALT199* in complex with 20-bp DNA (ipTM = 0.80; pTM = 0.84). Left, complexes displayed with surface electrostatics (−10 to +10, red to blue) in ChimeraX. Right, cartoon representations of the complexes displayed in ChimeraX. (**D**) Sequence alignment of DancE homologs. DancE was aligned with CD-NTase from *Escherichia coli KCJK1734*, *Pectobacterium parmentieri RNS08.42.1A* and *Alteromonas sp. ALT199* using Clustal W (*26*), and the alignment was visualized by ESPript (*27*).

**Fig. S6.**
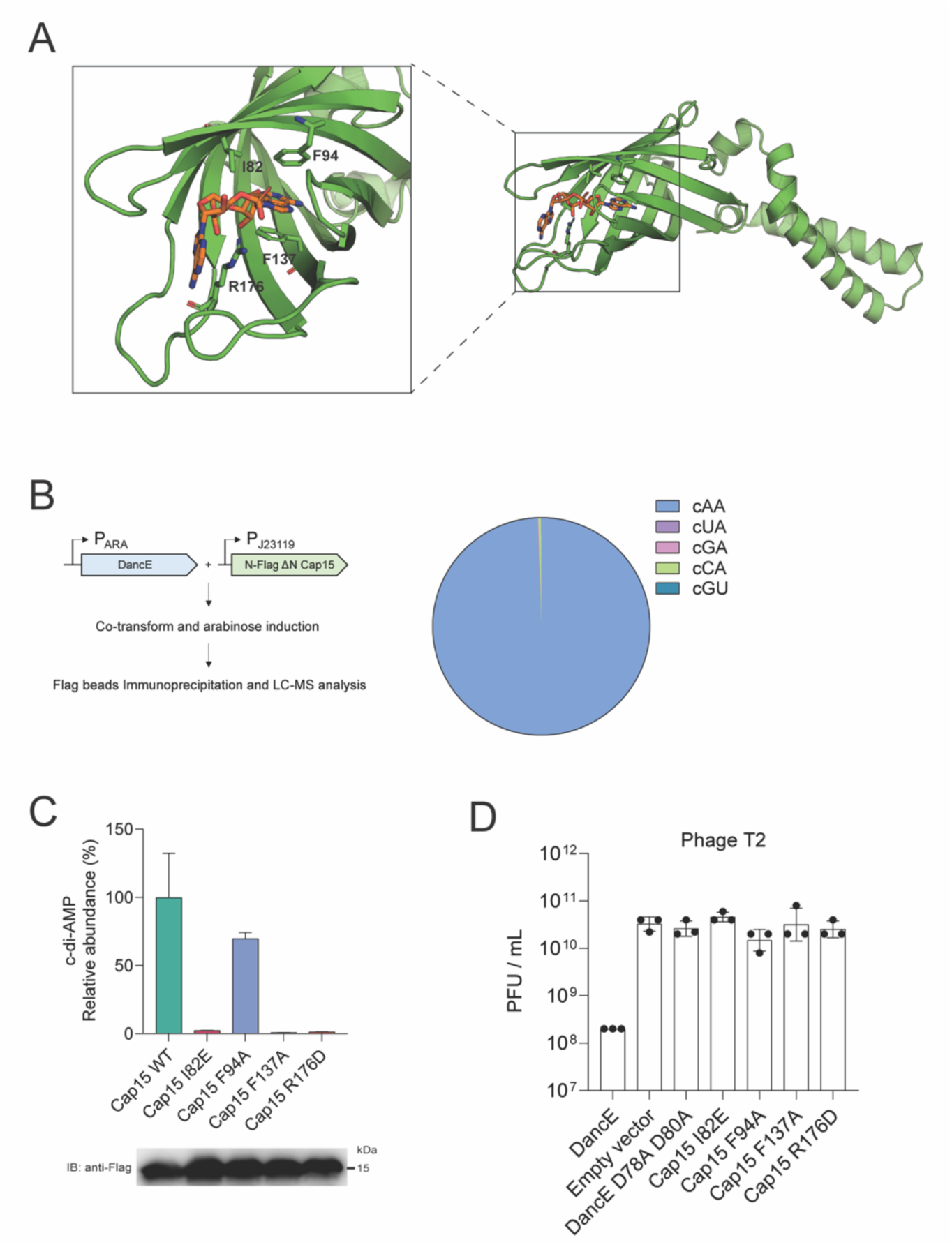
Cap15 from DancE CBASS binds c-di-AMP. (**A**) AlphaFold 3-predicted structure of the Cap15–c-di-AMP complex displayed in PyMOL (ipTM = 0.76; pTM = 0.70). (**B**) Left, schematic of the experimental workflow. Right, c-di-AMP (cAA) is the predominant *in vivo* product bound by Cap15. The data are representative of at least three independent experiments. cUA, c-UMA-AMP; cGA, c- GMP-AMP; cCA, c-CMP-AMP; cGU, c-GMP-UMP. (**C**) Mutations in the nucleotide-binding domain of truncated Cap15 (residues 69–195) impair c-di-AMP binding. Upper, relative abundance of c-di-AMP co-purified with wild-type (WT) Cap15 or the indicated Cap15 mutants (all truncated version). Data are mean ± s.d. from three independent replicates. Lower, *in vivo* expression of truncated Cap15 proteins. N-terminal Flag-tagged Cap15 constructs were expressed under the control of the J23119 promoter. Anti-Flag antibody (Sigma) was used for immunoblotting. (**D**) Efficiency of plating (EOP) of T2 phage on *E. coli* MG1655 expressing wild-type operon, empty vector (no operon), D78A D80A mutant (DancE catalytic dead) and indicated Cap15 mutants of DancE CBASS. Data are mean ± s.d. from three biological replicates.

**Fig. S7.**
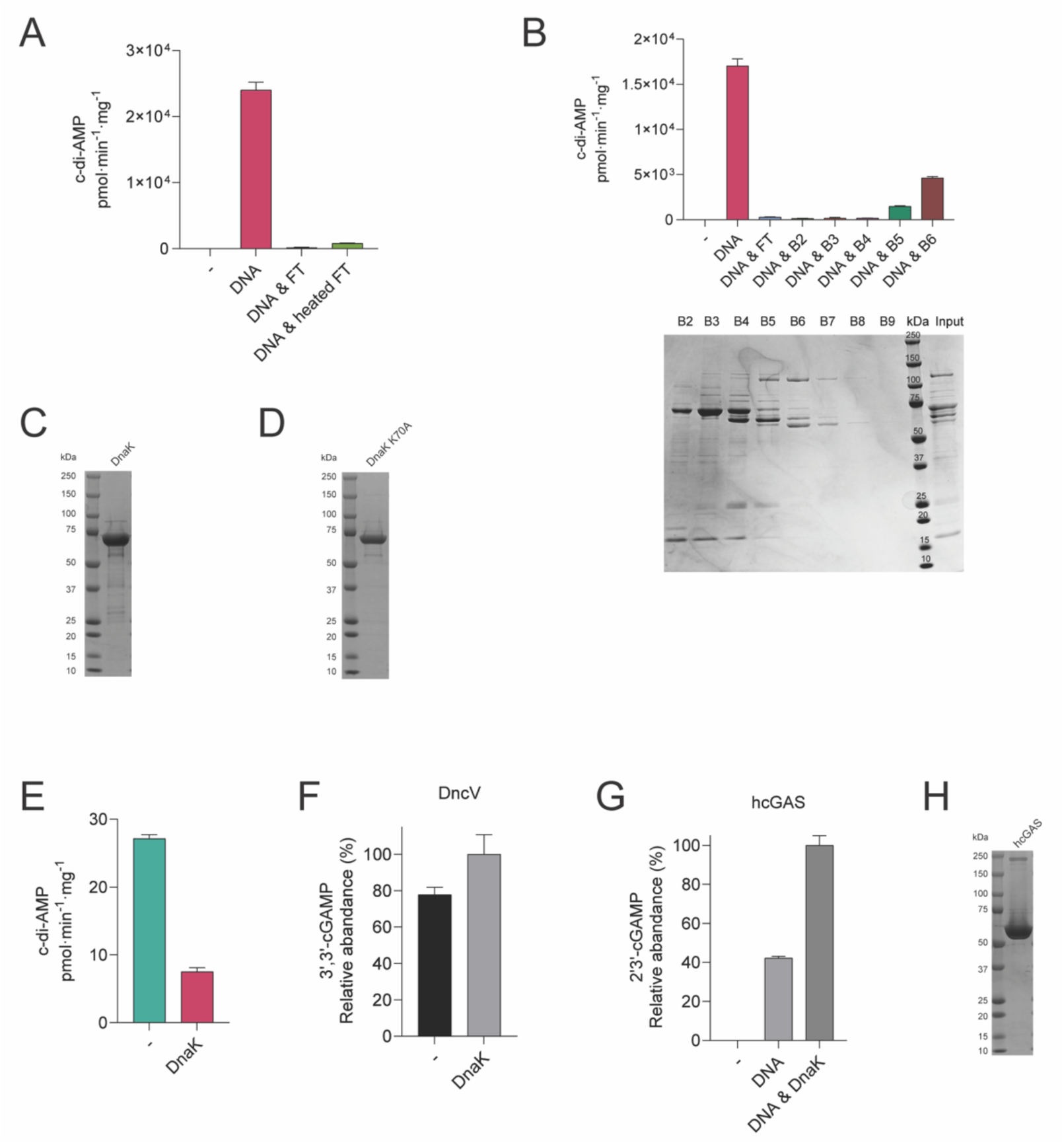
DnaK prevents DNA-dependent activation of DancE. (**A**) Wild-type DancE (0.8 µM) was incubated with or without *E. coli* genomic DNA (5 ng/µl) in the presence or absence of native or heat-treated (95 °C for 10 min) heparin column flow through (FT, 10% v/v) for 2 h. Data are mean ± s.d. from three independent replicates. (**B**) Upper, wild type DancE protein (0.8 µM) was incubated with or without *E. coli* genomic DNA (5 ng/µl) and heparin FT (5% v/v) or the indicated Q column fractions (B2–B6) derived from the heparin column FT for 2 h. Data are mean ± s.d. from three independent replicates. Lower, Coomassie blue-stained SDS– PAGE showing fractionation of the heparin column FT by anion-exchange chromatography. (**C**) Purified recombinant wild-type DnaK protein analyzed by SDS–PAGE and Coomassie blue staining. (**D**) Purified recombinant DnaK K70A mutant protein analyzed by SDS–PAGE followed by Coomassie blue staining. (**E**) Wild type DancE protein (2 µM) was incubated with or without DnaK (8 µM) for 16 h. Data are mean ± s.d. from three independent replicates. (**F**) Wild-type DncV protein (0.8 µM) was incubated with or without DnaK protein (8 µM) for 1 h. Data are mean ± s.d. from three independent replicates. (**G**) Wild-type human cGAS protein (0.8 µM) was incubated with or without plasmid (5 ng/µl) and DnaK protein (8 µM) for 1 h. Data are mean ± s.d. from three independent replicates. (**H**) Purified recombinant human cGAS protein analyzed by SDS–PAGE and Coomassie blue staining.

**Fig. S8.**
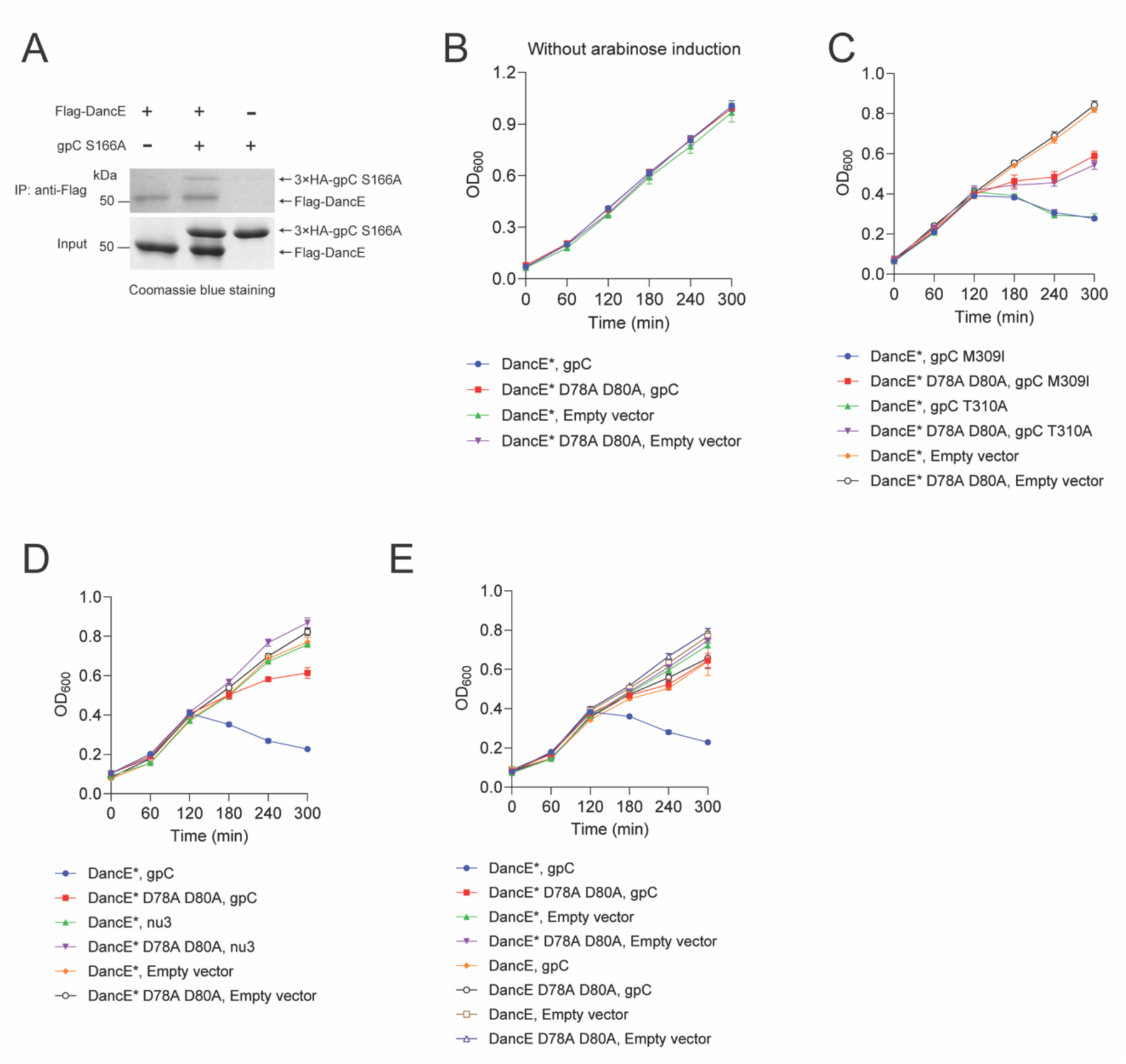
Lambda phage protease gpC triggers DancE activation. (**A**) Immunoprecipitation analysis of DancE-gpC interaction. Purified Flag–DancE wild-type and 3×HA gpC S166A proteins were incubated with anti-Flag M2 magnetic beads (Sigma) for Immunoprecipitation analysis. Eluted samples are analyzed by SDS–PAGE and Coomassie blue staining. (**B**)-(**E**), Reconstituted CBASS activation measured by cell growth curves. (**B**) Effects of wild-type gpC or empty vector on the growth of cells expressing DancE A33R (DancE*) or the catalytically inactive DancE* D78A D80A mutant, without arabinose induction. (**C**) Effects of gpC escaper mutants (M309I and T310A) or empty vector on the growth of cells expressing DancE* or DancE* D78A D80A mutant. Data for the empty vector group are shared with Fig. 5E. (**D**) Effects of wild-type gpC, wild-type Nu3, or empty vector on the growth of cells expressing DancE* or DancE* D78A D80A mutant. Data for the gpC and empty vector groups are shared with Fig. 5F. (**E**) Effects of wild-type gpC or empty vector on the growth of cells expressing DancE*, wild-type DancE, or the corresponding catalytically inactive D78A D80A mutants. Unless otherwise indicated, 0.25% (w/v) arabinose was added at 0 min to induce gpC or Nu3 expression. Data are mean ± s.d. from three biological replicates.

**Table S2.** Data collection and refinement statistics.

| **Data collection** |  |  |
| --- | --- | --- |
| Crystal | THFA-bound DancE<br>(PDB 37ZU ) | Apo DancE<br>(PDB 37ZT) |
| Space group | I432 | P6<sub>4</sub>22 |
| Cell constants (Å) | a = 252.69 | a = 152.44, c = 175.38 |
| Wavelength (Å) | 0.97946 | 1.00000 |
| Resolution range (Å) | 39.95 – 2.85<br>(2.90 – 2.85) | 38.11 – 2.55 (2.59 – 2.55) |
| Unique reflections | 31,914 (1,579) | 39,873 (1,935) |
| Multiplicity | 41.1 (31.1) | 27.0 (25.7) |
| Data completeness (%) | 100.0 (100.0) | 99.9 (100.0) |
| $R_{\text{merge}}$ (%)<sup>a</sup> | 22.0 (338.9) | 7.2 (275.1) |
| $R_{\text{pim}}$ (%)<sup>b</sup> | 3.4 (61.3) | 1.4 (55.3) |
| CC<sub>1/2</sub> (last resolution shell) | 0.499 | 0.556 |
| I/σ(I) | 22.8 (1.3) | 57.0 (1.3) |
| Wilson $B$-value (Å<sup>2</sup>) | 55.5 | 36.9 |
| **Refinement** |  |  |
| Resolution range (Å) | 39.95 – 2.86<br>(2.86 – 2.93) | 38.11 – 2.55 (2.61 – 2.55) |
| No. of reflections $R_{\text{work}}/R_{\text{free}}$ | 31,023/1,999 (1,352/93) | 38,472/2,000 (1,494/82) |
| Data completeness (%) | 97.21 (64.40) | 96.5 (56.7) |
| Atoms (non-H<br>protein/ligand/ions/solvent/waters) | 3,525/32/6/32/NA | 3,414/NA/1/8/134 |
| $R_{\text{work}}$ (%) | 19.2 (32.7) | 23.2 (31.4) |
| $R_{\text{free}}$ (%) | 20.8 (38.1) | 27.5 (36.8) |
| R.m.s.d. bond length (Å) | 0.006 | 0.002 |
| R.m.s.d. bond angle (°) | 0.803 | 0.492 |
| Mean $B$-value (Å<sup>2</sup>)<br>(protein/ligands/ions/solvent/waters) | 55.67/76.99/66.72/63.50/<br>NA | 44.54/NA/38.38/61.44/36.66 |
| Ramachandran plot (%)<br>(favored/additional/disallowed)<sup>c</sup> | 96.8/3.2/0.0 | 97.9/2.1/0.0 |
| Clashscore/Overall score<sup>c</sup> | 1.82/1.22 | 1.18/0.86 |
| Maximum likelihood coordinate error | 0.31 | 0.23 |
| Missing residues | 306-311. | 306-311, 432-442. |

Data for the outermost shell are given in parentheses.
<sup>a</sup> $R_{\text{merge}} = 100 \sum_h \sum_i |I_{h,i} - \langle I_h \rangle| / \sum_h \sum_i \langle I_{h,i} \rangle$ , where the outer sum (h) is over the unique reflections and the inner sum (i) is over the set of independent observations of each unique reflection.
<sup>b</sup> $R_{\text{pim}} = 100 \sum_h \sum_i [1/(n_h - 1)]^{1/2} |I_{h,i} - \langle I_h \rangle| / \sum_h \sum_i \langle I_{h,i} \rangle$ , where $n_h$ is the number of observations of reflections **h**.
<sup>c</sup>As defined by the validation suite MolProbity (Chen, V.B., Arendall, W.B.A., Headd, J.J., Keedy, D.A., Immormino, R.M., Kapral, G.J., Murray, L.W., Richardson, J.S., Richardson, D.C. (2010) *MolProbity*: all-atom structure validation for macromolecular crystallography. *Acta Cryst.* **D66**, 12-21.).

